# GREMLIN1 disrupts intestinal epithelial-mesenchymal crosstalk to induce a wnt-dependent ectopic stem cell niche via stromal remodelling

**DOI:** 10.1101/2024.04.28.591245

**Authors:** EJ. Mulholland, HL. Belnoue-Davis, GN. Valbuena, N. Gunduz, A. Ligezal, S. Biswasl, E. Gil Vasquez, S. Omwengal, N. Nasreddinl, M. Hodder, LM. Wang, S. Irshad, AS. Ng, LK. Jennings, KS. Midwood, N. Dedi, R. Ridgeway, T. Phesse, JE. East, IPM. Tomlinson, GCG. Davies, O. Sansom, SJ. Leedham

**Author notes:** **Correspondence:** Address correspondence to, Simon Leedham, Wellcome Trust Centre for Human Genetics, Roosevelt Drive, Oxford, OX3 7BN, UK. Joint first authors (EJ. Mulholland, HL. Belnoue-Davis).

## Abstract

In homeostasis, counterbalanced morphogen signalling gradients along the vertical axis of the intestinal mucosa regulate the fate and function of epithelial and stromal cell compartments. Here, we used a disease-positioned mouse, and human tissue, to explore the consequences of pathological Bone Morphogenetic Protein (BMP) signalling dysregulation on epithelial-mesenchymal interaction. Aberrant pan-epithelial expression of the secreted BMP antagonist GREM1, resulted in ectopic crypt formation with lineage tracing demonstrating the presence of *Lgr5(-)* stem/progenitor cells. Isolated epithelial cell *Grem1* expression had no effect on individual cell fate, indicating an intercompartmental impact of mucosal-wide BMP antagonism. Treatment with a novel anti-Grem1 antibody abrogated the polyposis phenotype, and triangulation of specific pathway inhibitors defined a pathological sequence of events, with wnt-ligand dependent ectopic stem cell niches formed through stromal remodelling following BMP disruption. These data support an emerging co-evolutionary model of intestinal cell compartmentalisation based on bidirectional regulation of epithelial-mesenchymal cell fate and function.

**One Sentence Summary:** Pathological epithelial GREM1 expression induces therapeutically reversible ectopic stem cell niches through stromal remodelling

## Introduction

The unique, crypt-based architecture of the gut mucosa facilitates the study of intestinal cell fate. Each crypt is supported by a discrete population of adult intestinal stem cells, and luminally-directed daughter cell migration results in continual epithelial turnover along a crypt-to-villus cell escalator ^1^. This stem-to-differentiated functional segregation results from the establishment of opposing gradients of intercompartmental morphogenetic signalling ^2,3^. At the luminal surface, stromally secreted Bone Morphogenetic Protein (BMP) ligands mediate cell differentiation and apoptosis, whereas in the crypt base, restricted Wnt and Notch signalling drives epithelial stemness, transit amplifying cell division and regulates progenitor cell fate. Exclusive sub and peri-crypt stromal cell expression of secreted, ligand-sequestering BMP antagonists (BMPi) within the stem cell niche, protects against premature differentiation of crypt base columnar stem cell populations^4–6^. Developmentally, these polarised signalling gradients are both a cause and consequence of the distinct architecture of the intestinal mucosa, and the inter-relationship of the epithelial and stromal cell compartments. In the developing small intestine, villus formation precedes crypt morphogenesis. Physical buckling of the epithelial surface generates projecting villi that results in the formation of a sub-epithelial, BMP-expressing stromal cell aggregation, known as the villus cluster ^7^. This pro-differentiation signalling at the luminal surface repels putative stem cells to the base of the villus where developing crypts generate restricted, wnt-high crypt basal stem cell niches ^7^. The impact of polarised signalling gradients in regulating homeostatic epithelial cell fate is established but recent findings also demonstrate the morphogenetic regulation of stromal cell function along the crypt-villus axis. Kraiczy *et al* used cell marker expression to segregate intestinal fibroblast cell populations into distinct functional subsets^8^. Mesenchymal expression of stem cell supporting signalling factors, such as R-Spondins and BMPi, were restricted to subcryptal CD81(+) PDGFRA(lo) trophocytes, and pericryptal CD81(-) CD55(+) PDGFRA(lo) stromal cells, resulting in the maintenance of a discrete crypt basal stem cell niche capable of supporting stem cells. The authors proposed that BMP ligand signalling arising from luminal aggregates of PDGFRA(hi) sub-epithelial myofibroblasts (SEMF) regulates this variable stromal cell activity, suppressing stem cell niche supporting functionality at the luminal surface ^8^. Consequently intercompartmental secreted BMP signalling, generates a polarised signalling gradient along the intestinal vertical axis, that regulates not only epithelial cell fate but also stromal cell function to restrict appropriate stem cell niche activity exclusively to the crypt base.

This interdependence between mucosal structure and function regulated by secreted signalling pathways, renders the intestine susceptible to disruption of the morphogen gradient balance. The majority of intestinal lesions arise through epithelial mutation-induced activation of the wnt pathway, however pathological inactivation of BMP signalling is also capable of initiating human polyp formation. This is epitomised by the human polyposis syndromes Juvenile Polyposis and Hereditary Mixed Polyposis Syndrome (HMPS), that arise through germline mutations in the BMP pathway. In HMPS, a duplication mutation on chromosome 15 causes ectopic pan-epithelial expression of the secreted BMPi, *GREMLIN1*, which is normally restricted to exclusive expression by sub and peri crypt stromal cells ^9^. In mouse models, forced epithelial expression of secreted BMPi, such as *Noggin* or *Grem1* leads to architectural distortion with the generation of ectopic crypts ^10–12^. These ectopic crypt foci (ECF) are populated by Lgr5(-) progenitor cells, that proliferate and acquire oncogenic driver mutations, with eventual tumorigenesis arising from a cell-of-origin situated outside of the regulatory confines of the crypt base ^12^.

Here, we use a disease-positioned animal model of HMPS to mechanistically explore the consequences of pathological morphogen gradient dysregulation on epithelial and mesenchymal cell compartment structure and function. We investigate the establishment and regulation of ectopic stem cell niches and demonstrate the impact of novel therapeutic intervention on the intestinal neoplasia initiated by disruption of polarised BMP signalling gradients.

## Results

### Lineage tracing from ectopic crypt cells

Our previous work had generated a novel disease-positioned mouse model of Hereditary Mixed Polyposis Syndrome (HMPS) called *Vil1-Grem1* using the *Villin1* promoter to drive aberrant intestinal epithelial expression of the secreted BMP antagonist *Grem1*. This induces a pronounced pan-intestinal polyposis with lesions that phenocopy the mixed polyp histology of the human condition, including the formation of ectopic crypts, best seen within the villus in small intestinal lesions ^12^. We have previously shown aberrant cell proliferation but notable lack of expression of the crypt base columnar stem cell marker *Lgr5* in villus ectopic crypt cells ^12^ and through further analysis of ectopic crypt cell expression, we were unable to identify any unique and restricted ectopic crypt stem cell markers. However, we did identify a number of shared markers that were expressed in both the homeostatic progenitor cell population in the normal crypt base as well as in the pathological progenitor cells situated in the ectopic crypt (e.g. *SOX9*) (Figure 1A). In order to functionally demonstrate the stem cell potential of ectopic crypt foci, we generated *Sox9-CreER^T^*^2^*; Rosa26^YFP^; Vil1-Grem1* mice to assess whether ectopic crypt SOX9(+) cells were capable of generating lineage tracing ribbons *in vivo*. We aged animals for 120 days to establish a pan-intestinal polyposis phenotype and then induced chimeric recombination with low dose tamoxifen to activate YFP expression in a small number of SOX9(+) cells (Figure 1B). Recombined animals were sacrificed after 30 days and chromogenic immunohistochemistry against YFP was used to identify lineage traced cell populations arising from *Sox9*+ve cells situated within both the normal and ectopic crypt bases. Although, we saw frequent lineage tracing ribbons arising above the crypt isthmus in small intestinal polyps, we recognised that in two dimensional sections, these visualised traced cells could conceivably have arisen from proximally situated *Sox9+* cell recombination, within the normal crypt base. To exclude this, we undertook whole lesion serial sectioning and used tissue alignment and digital reconstruction of individual polyps (HeteroGenius, Leeds, UK) to examine the three-dimensional path of lineage traced cells (Supplementary movies 1,2). This allowed us to map tracing ribbons in their entirety and track the path of YFP labelled cell progeny (Figure S1). Using this technique we were able to identify tracing ribbons, originating above the crypt base from villus ectopic crypts, that were spatially distinct from traces emerging from neighbouring crypt basal cells (Figure 1C). Together this work demonstrated that proliferating *Sox9*+ve cells situated in the ectopic crypts within the villus of *Vil1-Grem1* mice are capable of generating multicellular lineage tracing ribbons, and thus show functional stem cell potential *in vivo*.

**Figure 1.**
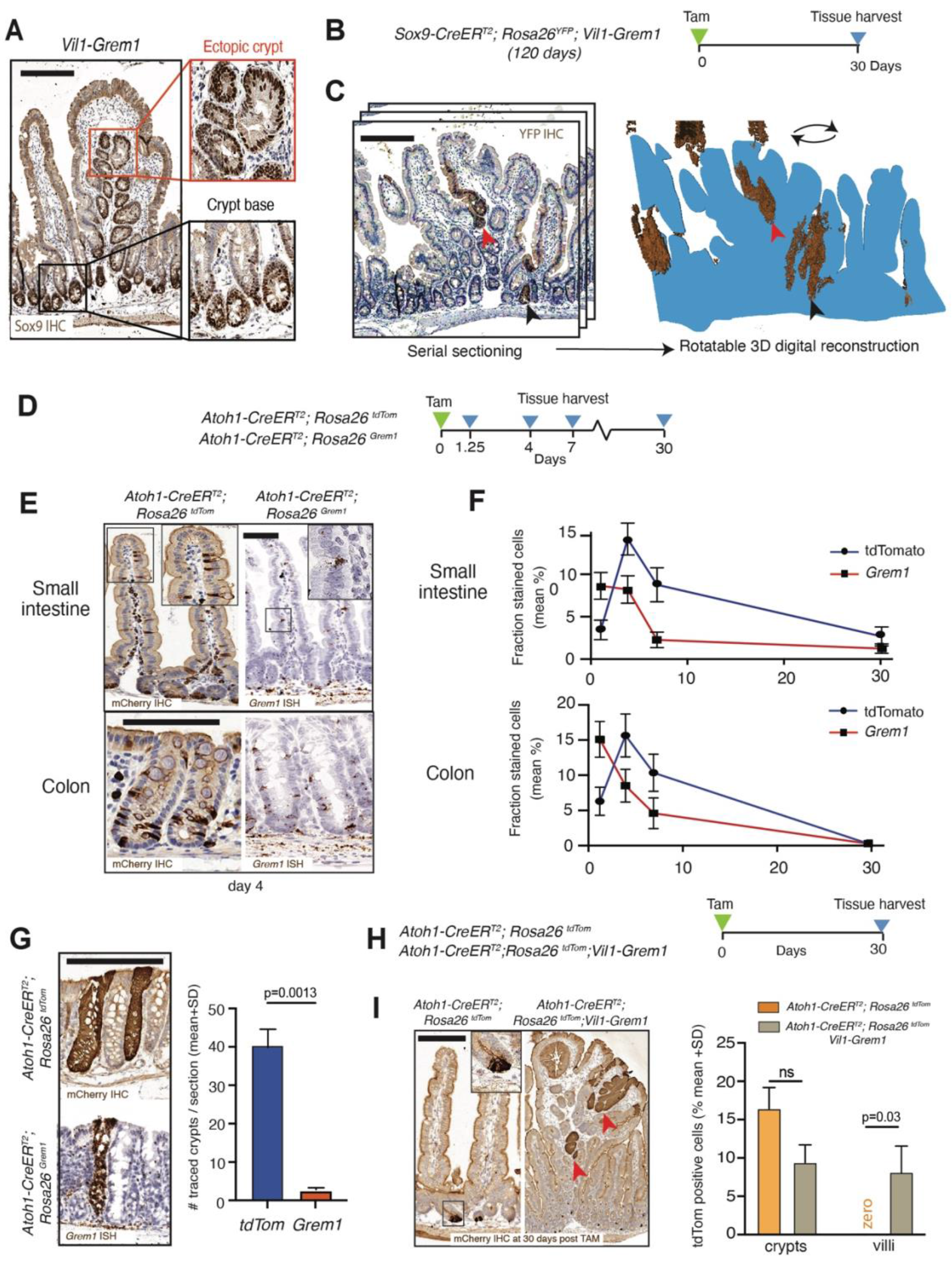
Ectopic crypt lineage tracing and secretory cell fate. **A.** SOX9 immunohistochemical staining in 120 day *Vil1-Grem1* polyp shows specific nuclear staining in normal crypt base (black box) and ectopic crypt (red box) **B.** Schematic shows recombination and harvesting timepoint for *Sox9-CreER^T2^; Rosa26^YFP^;Vil1-Grem1* mouse model. **C.** Tissue serial sectioning and lineage tracing ribbon identification with anti-YFP immunohistochemistry (brown), followed by tissue alignment and polyp 3D reconstruction of lineage tracing ribbons (brown) (aligns with Figure S1). **D.** Schematic showing recombination and harvesting of secretory cell mouse models. **E.** Secretory cell staining in small intestine and colon 4 days post-recombination in *Atoh1-CreER^T2^,Rosa26^tdTom^*or *Atoh1-CreER^T2^; Rosa26^Grem1^* mice stained with mCherry IHC or *Grem1* ISH respectively. **F.** Number of mCherry IHC (blue lines) or *Grem1* ISH (red lines) stained small intestinal or colonic epithelial cells over time following recombination (n=5 mice per group) **G.** Quantification of number of fully lineage traced colonic crypts across gut roll sections at day 30 post recombination in *Atoh1-CreER^T2^,Rosa26^tdTom^* or *Atoh1-CreER^T2^; Rosa26^Grem1^* mice stained with mCherry IHC or *Grem1* ISH respectively (n=5 mice per genotype, t test, p value as stated). **H.** Schematic shows recombination and harvesting of secretory mouse models in a *Vil1-Grem1* disease setting**. I.** Staining and quantification of mCherry stained cells crypts and polyps in *Atoh1-CreER^T2^; Rosa26^tdTom^*and A*toh1-CreER^T2^; Rosa26^tdTom^; Vil1-Grem1* animals respectively at 30 days post recombination (n=5 per group, 2-way ANOVA, p-value as stated, ns=p>0.05). Scale bar 200µm throughout.

### Aberrant *Grem1* expression in individual secretory cells does not alter cell fate trajectory

In homeostasis, *Grem1* expression from sub and peri-crypt stromal cells is important for maintaining crypt basal *Lgr5(+)* stem cell number^5^, however the cell-intrinsic impact of aberrant *Grem1* expression on the fate of epithelial cells situated outside of the crypt basal niche is unknown. To assess this, we used *Atoh1-CreER^T^*^2^ to induce expression of either tdTomato marker (*Atoh1-CreER^T2^;Rosa26^tdTom^*) or *Grem1* (*Atoh1-CreER^T2^;Rosa26^Grem1^*) in individual ATOH1+ve secretory cell progenitors in steady state conditions (Figure 1D). We then tracked secretory cell fate through expression of either the active morphogen *Grem1* (with ISH), or the functionally neutral tdTomato marker (with IHC), taking care to quantify and contrast spatio-temporal fate of cells within animal groups when utilising these different methodological cell marking techniques (Figure 1E). In the small intestine, expression of neutral tdTomato marker was seen in crypt basal cells within 30 hours and was retained in small numbers of long-lived Paneth cells for 30 days. This pattern of Paneth cell fate trajectory was mirrored by individual cells expressing autocrine *Grem1*. Other tdTomato or *Grem1* expressing cells differentiated into goblet cells, moved along the crypt-villus axis and were appropriately lost over time through cell shedding. This loss was initially accelerated in individual cells aberrantly expressing *Grem1* (Figure 1F). In the colon, occasional crypt lineage tracing could be seen from *Atoh1* derived cells, as a consequence of cell de-differentiation to Lgr5(+) crypt-base-columnar cells ^13^ however, consistent with accelerated cell loss, the number of lineage traced crypts was significantly reduced in cells expressing autocrine *Grem1* (Figure 1G). These data show that in steady state conditions, individual cell-autonomous *Grem1* expression does not intrinsically promote epithelial cell de-differentiation, enhance functional stemness or prevent appropriate terminal differentiation of that cell.

Next, we generated *Atoh1-CreER^T2^; Rosa26^tdTom^; Vil1-Grem1* animals to compare the impact of pan-epithelial *Grem1* expression on secretory cell fate (Figure 1H). In contrast with normal small bowel, numerous confluent clones of TdTomato labelled cells were seen 30 days after recombination emerging from the ectopic crypts of polyps in *Atoh1-CreER^T2^; Rosa26^tdTom^; Vil1-Grem1* animals (Figure 1I). Thus, in contrast to single cell expression, pan-epithelial upregulation of BMPi leads to disrupted secretory cell fate with persistence of Atoh1 traced cell progeny within the pathologically altered villus compartment. Together these data show that isolated, single cell *Grem1* expression does not intrinsically alter fate through individual cell autocrine BMP antagonism. In contrast, pan-epithelial expression of *Grem1*, with concomitant disruption of mucosal-wide BMP signalling gradients does lead to the disruption of normal secretory cell fate determination and the emergence of an ectopic crypt stem population that is not driven exclusively by aberrant epithelial cell-intrinsic BMPi expression.

### Phenotype reversal through *Grem1* inhibition

The pathognomic mixed polyposis phenotype in HMPS^9,14^ and the *Vil1-Grem1* model is caused by aberrant epithelial expression of *Gremlin1*^12^. To see if we could reverse the effects of this secreted antagonist, we undertook short and long-term treatment of *Vil1-Grem1* animals with a novel anti- Grem1 antibody (UCB Ab7326 mIgG, UCB Pharma) which blocks sequestering of BMP ligands ^15^, commencing treatment either at weaning (35 days old) or in animals with an established polyposis (120 days old) (Figure 2A). Animals treated with UCB Ab7326 showed a remarkable reversion of the characteristic *Vil1-Grem1* pan-intestinal polyposis phenotype (Figure 2B and S2). Treatment profoundly increased mean mouse survival, with long-term treated mice succumbing to old age, rather than burden of intestinal disease (Figure 2C). Temporally spaced sampling in mice with established disease allowed assessment of the timing of antibody impact on established mouse polyposis phenotype. Macroscopically, 4 weeks of antibody treatment significantly reduced the size of the pathologically enlarged and thickened intestinal mucosa seen in these animals (Figure 2B,D). Microscopically, there was re-imposition of recognisable villus architecture, with restoration of normal villus patterns of the CK20 differentiation cell marker. UCB Ab7326 antibody treatment led to a rapid and profound abrogation of polyposis, near elimination of villus ectopic crypts and absence of villus proliferating KI67(+), SOX9(+) cells. OLFM4, EPHB2 and lysozyme expressing cells were once again appropriately confined to the base of the intestinal crypts in treated animals (Figure 2D,E, Figure S2). These data show that aberrant epithelial *Grem1* expression induces reversible intestinal architectural change and that the mixed polyposis phenotype can be both prevented and reversed through sequestering inhibition of Grem1.

**Figure 2.**
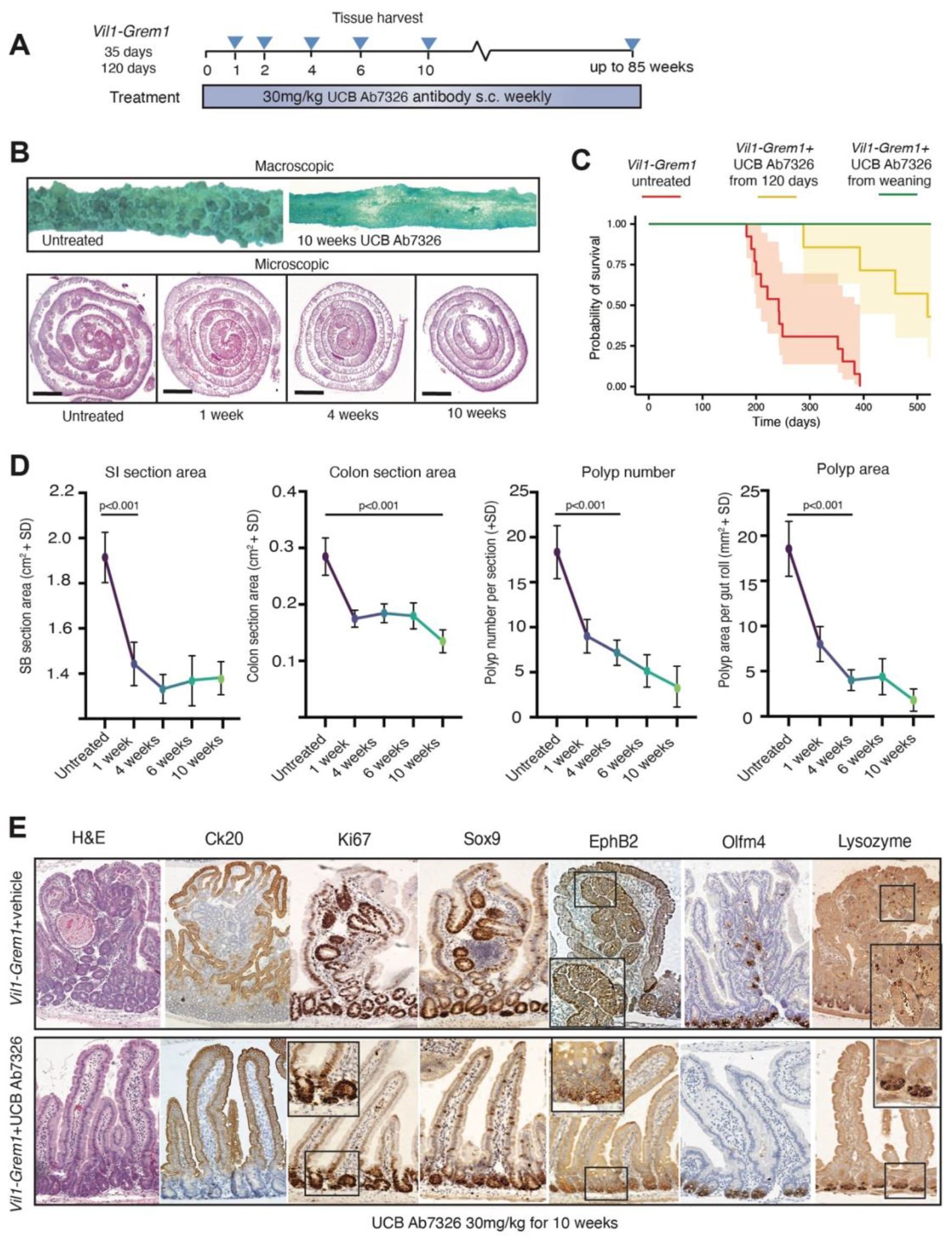
Phenotype reversal through UCB Ab7326 treatment. **A.** Schematic shows treatment schedule and harvesting timepoints of *Vil1-Grem1* mice treated with UCB Ab7326 . **B.** Representative macroscopic and H&E images of proximal small intestinal sections harvested from untreated and UCB Ab7326 treated *Vil1-Grem1* mice over a range of timepoints. (scale bars 0.25cm) **C.** Kaplan-Meier Survival Curve showing impact on survival of UCB Ab7326 initiated at weaning, (green line, n=11 mice, P=1.5E-06) or from 120 days (yellow line, n=7 mice, p=0.00421) in comparison to untreated animals (n=13 mice, red line). Significance calculated using log-rank tests, with the Benjamini-Hochberg correction for multiple testing. **D.** Quantification of *Vil1-Grem1* small bowel (SB) and colonic section surface area, polyp number and polyp area following variable time treatment with UCB Ab7326 antibody. (n=6 mice per group, one way ANOVA with Dunnett post-hoc corrections, p values as stated). **E.** Immunohistochemical analysis to show reversion of *Vil1-Grem1* ectopic crypt phenotype and restoration of normal crypt-villus staining patterns of KI67, SOX9, EPHB2, CK20 and Lysozyme following 10 weeks UCB Ab7326 therapy. Scale bar 200µm.

### Aberrant pan-epithelial Grem1 expression causes villus intercompartmental remodelling with de-repression of stem cell supporting fibroblast cells

In the normal intestine, Kraiczy et al have recently shown that proximity to luminal BMP ligand source represses stem cell niche fibroblast cell functionality, whereas at the crypt base distance from ligand source and the expression of *Grem1*, relieves this repression, favouring stem cell supporting stromal cell phenotypes^8^. In light of this, and the lack of impact of autocrine *Grem1* expression on cell-intrinsic epithelial cell fate (Figure 1E,F), we reasoned that pathological pan-epithelial expression of secreted *Grem1* in *Vil1-Grem1* animals could have an intercompartmental impact on underlying cell populations. Furthermore, the success of UCB Ab7326 treatment in phenotypic reversal provided a tool to assess the impact of partial restoration of polarised BMP signalling on different cell compartments. To investigate this, we used a variety of technologies to spatially assess the epithelial, stromal, matrix and immune compartments in UCB Ab7326 treated and untreated *Vil1-Grem1* mice.

To assess the distribution of BMP pathway activity along the crypt-villus axis, we used chromogenic pSMAD1,5 staining. In untreated animals, aberrant epithelial Grem1 secretion not only abrogated epithelial pSMAD1,5 staining (Figure 3A) but also altered the proportion, distribution and intensity of fibroblast cell expression (Figure 3B), indicating disruption of both epithelial and stromal BMP signalling gradients. Treatment with UCB Ab7326 antibody did not eliminate epithelial autocrine Grem1 function and restore epithelial homeostatic BMP signalling gradients - marked by ongoing loss of epithelial pSMAD1,5 staining (Figure 3A). In contrast, antibody treatment did mostly revert villus stromal BMP signalling, with re-emergence of crypt isthmus and villus fibroblast pSMAD1,5 staining (Figure 3B).

**Figure 3.**
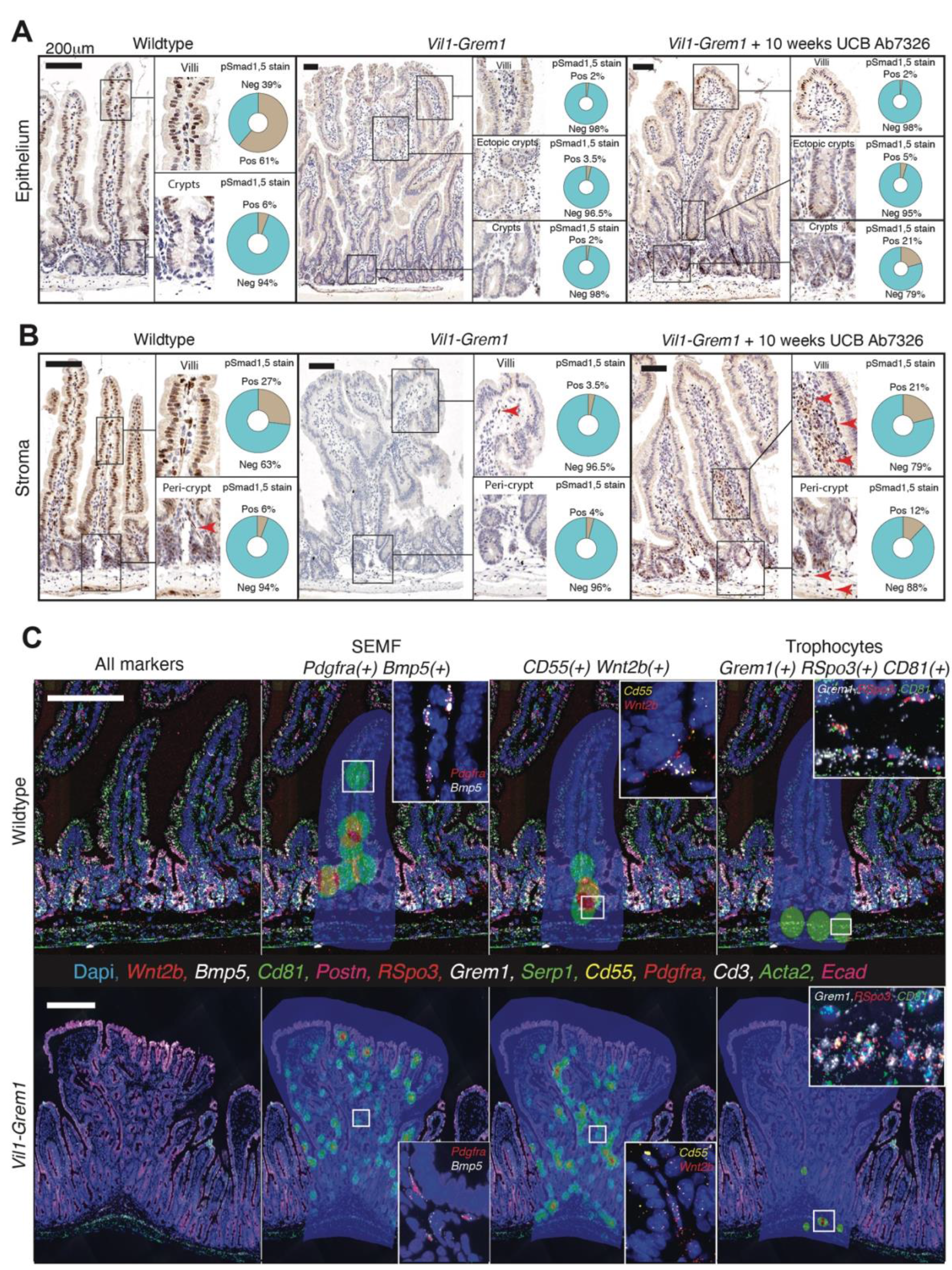
Aberrant pan-epithelial *Grem1* expression causes intercompartmental remodelling with de-repression of stem cell supporting fibroblast cells. **A.** pSMAD1-5 immunohistochemistry shows physiological villus epithelial cell nuclear staining from the crypt isthmus upwards (brown stain). Pan epithelial expression loss is seen in *Vil1-Grem1* mice, with little reversion of epithelial stain seen following 10 weeks UCB Ab7326 therapy. Automated stain quantification using digital pathology platform used to exclude stromal cell staining (QuPath) (n=3 mice per group, >15 villi or polyps total). **B.** pSMAD1-5 immunohistochemistry shows physiological stromal cell staining in the villus stomal compartment (brown stain). Loss of expression is seen in untreated *Vil1-Grem1* animals as a consequence of intercompartmental BMP antagonism. Treatment with UCB Ab7326 recovers stromal expression levels of pSMAD1-5 in treated *Vil1-Grem1* mice. Automated stain quantification using digital pathology platform used to exclude epithelial cell staining (QuPath, n=3 mice per group, >15 villi or polyps total). **C.** Hi-Plex *in situ* hybridisation (12 markers) was used to identify and topographically map functionally distinct fibroblast subtypes (SEMF, *Cd55(+) Wnt2b(+),* and Trophocytes) in wild-type and *Vil1-Grem1* mouse tissue (n=5 mice per group). Image analysis software was used to identify each subtype and map cell location back onto the tissue based on density distribution (coloured circles, within shaded areas - HALO, Indica Labs). Scale bars 200µm.

Given the impact of epithelial *Grem1* expression on the underlying stroma, we generated a custom Hi-Plex *ISH* panel to explore the effect of disrupted BMP gradients on the spatial distribution of different homeostatic functional stromal cell phenotypes ^6,8^ (Figure S3A). In wild-type animals we were able to confirm the crypt isthmus and villus distribution of sub-epthelial myofibroblasts (SEMF), the restricted peri-cryptal distribution of *Cd55*(+) *Wnt2b*(+) cells and the sub-cryptal distribution of trophocytes. However, in *Vil1-Grem1* animals, the crypt-restriction of *Cd55(+) Wnt2b(+)* cell populations was lost, with a marked expansion of this cell type up into the villus compartment, often surrounding ectopic crypt cells in the middle of the polyp (Figure 3C and S3B).

Consistent with this remodelled stromal cell landscape, we also saw significant differences in the proportion and distribution of extracellular matrix proteins in *Vil1-Grem1* animals, with an expansion of COL1A1, MMP-3 and LAMININ expression in the polyps (Figure S3C). Multiplex staining of key innate and adaptive immune cell populations revealed a significant influx of T-helper and T-reg cells and a reduction of T-cytotoxic cells in polyps (Figure S3D). Previous work has shown that *Grem1* expressing fibroblastic reticular cells in secondary lymphoid structures support dendritic cell function and provide a niche for CD4(+) T-Cells^16^, although the functional role of *Grem1* expression in this process was not directly tested. These data provide circumstantial evidence of a possible role for homeostatic BMP signalling in maintaining normal T-cell immunity, however further study on mechanisms of immune cell regulation would be needed and are beyond the scope of this study.

Together, this work shows that disruption of homeostatic stromal BMP signalling gradients through epithelial *Grem1* expression has an impact on sub-epithelial fibroblast populations causing a de-repression of *Cd55(+) Wnt2b(+)* fibroblasts, which then aberrantly expand beyond the normal spatial confines of the peri-crypt region.

### Formation of the ectopic stem cell niche requires ligand-dependent Wnt signalling

Following our demonstration of BMPi-dependent remodelling of the villus stromal cell compartment, we wished to explore the signalling pathway constituents of the ectopic stem cell niche. To do this we combined IHC and *in situ hybridisation* to identify morphogens expressed in ectopic crypts and their supporting stromal cells.

First, we spatially assessed the activity of the Wnt pathway in ectopic crypts at ligand, receptor and target gene level. In comparison with normal small intestinal tissue, *in situ* hybridisation of key intestinal wnt ligands (Figure 4A and S4A) showed aberrant distribution and increased expression of *Wnt2b* predominantly around the ectopic crypts with some, but not all, of this ligand coming from co-stained *Cd55(+)* fibroblasts (Figure 4A,E). Increased non-canonical *Wnt5a* ligand expression was seen, arising from a different stromal cell population, co-stained with periostin and situated peripherally around the polyp edge (Figure 4A and S4B). At a receptor level, both *Fzd5* and *Lgr4* were heavily upregulated in the epithelium of *Vil1-Grem1* mice, especially in ectopic crypt cells (Figure 4A). From a target gene perspective, both SOX9 (IHC) and *Axin2(ISH)* were specifically aberrantly upregulated in proliferating ectopic crypts within the villus compartment (Figure 1A, 4A). Following UCB Ab7326 treatment, and the restoration of crypt/villus architecture, we saw reversion of most Wnt pathway activity markers to appropriate restricted expression in the crypts/isthmus.

**Figure 4.**
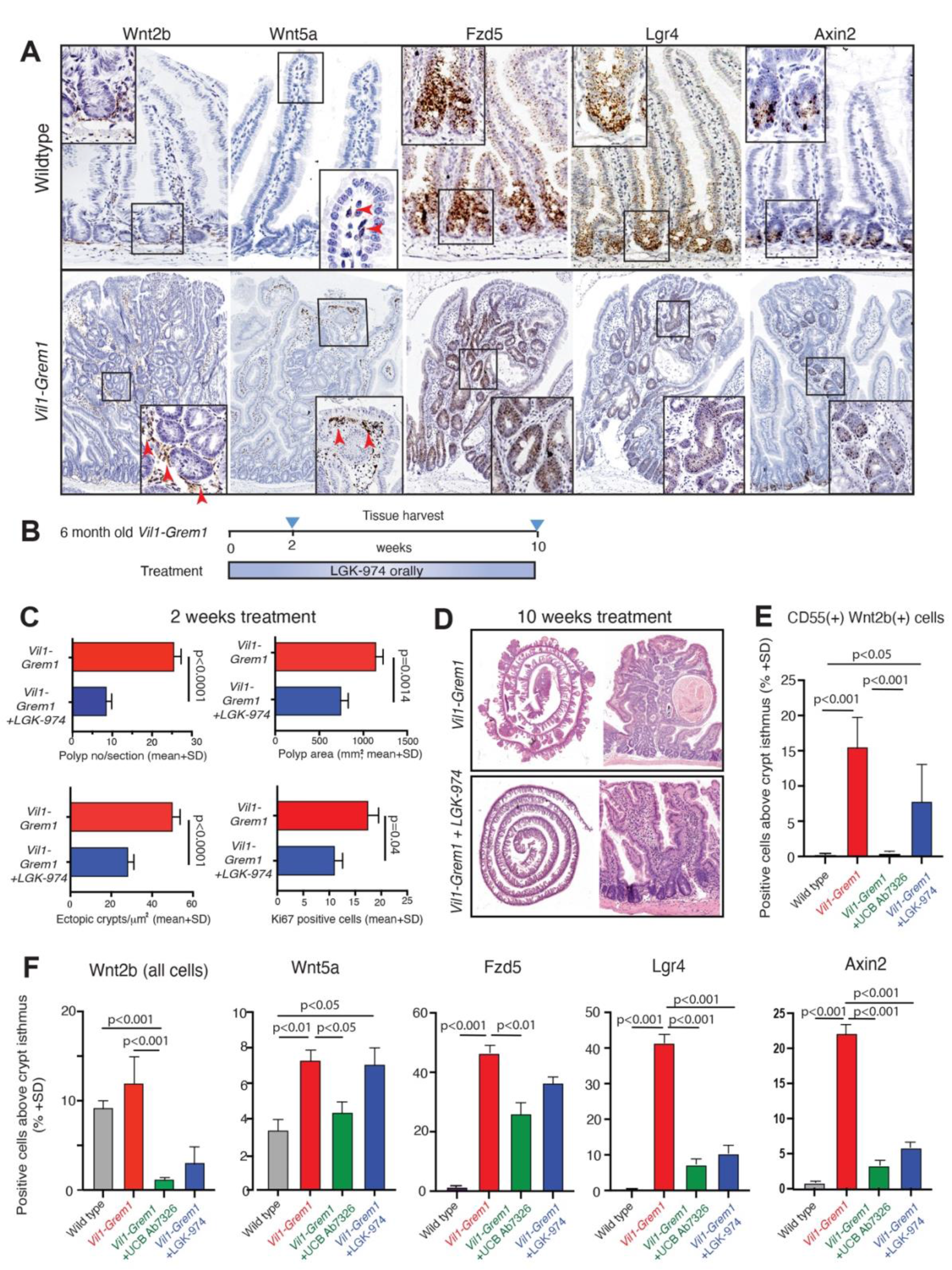
Formation of the ectopic niche requires ligand-dependent Wnt signalling. **A.** Representative images of *in situ* hybridisation of Wnt ligands (*Wnt2b, Wnt5a*), receptors (*Fzd5, Lgr4*) and target gene (*Axin2*) in *Vil1-Grem1* versus wild-type (C57BL/6J) small intestine. Scale bar 200µm **B.** Schematic shows treatment and harvesting timepoints of *Vil1-Grem1* animals treated with the porcupine inhibitor, LGK-974. **C.** Digital pathology-based quantification of: polyp number per section, polyp area, ectopic crypts and villus KI67+ cell number per gut roll following 2 weeks of porcupine inhibition (n=5 mice per group, t test, p values as stated). **D.** Representative H&E images of untreated and LGK-974 treated *Vil1-Grem1* mouse small intestine (10 weeks) showing loss of polyps and ectopic crypts but ongoing villus widening/deformity in treated animals. Scale bar 0.25cm for gut rolls **E.** Quantification of *Cd55(+) Wnt2b(+)* (ISH) cell proportion above the crypt isthmus in wildtype and *Vil1- Grem1* animals including after UCB Ab7326 and LGK-974 therapy (n=5 mice, t test, p values as stated) **F.** Quantification of wnt target, receptor and target stained cell proportion above the crypt isthmus in wildtype and *Vil1-Grem1* animals including after UCB Ab7326 and LGK-974 therapy (n=5 mice per group, t test, p values as stated)

Given this evidence for significant aberrant Wnt pathway activity in ectopic crypts we undertook pharmacological and genetic manipulation of the pathway. Firstly, we used a specific Porcupine inhibitor LGK-974 to inhibit palmitoylation and secretion of Wnt ligands in 120 day old *Vil1- Grem1* animals. Both short (2 week) and long-term (10 week) inhibition of wnt ligand secretion led to a significant reduction in polyp number, size, ectopic crypt formation and aberrant villus cell proliferation (Figure 4C,D). Porcupine inhibition did reduce abnormal villus compartment wnt activity and abrogate ectopic crypt formation, however unlike UCB Ab7326 antibody therapy, wnt ligand suppression did not restore macroscopically normal small intestinal architecture, with ongoing villus enlargement and deformity seen throughout the small bowel (Figure 4D). To interrogate this ongoing macroscopic change, we assessed key stromal cell populations in treated mice. Following UCB Ab7326 treatment we had noted a complete re-suppression of villus *Cd55(+) Wnt2b(+)* cells, however this was not the case after porcupine treatment where we saw ongoing disruption of stromal cell functional architecture and continuing presence of de-repressed *Cd55(+) Wnt2b(+)* cells in the villus compartment (Figure 4E).

Next, to interrogate signal transduction of ligand-dependent wnt signalling, we generated *Villin-CreER^T2^; Fzd5^fl/fl^; Vil1-Grem1* and *Villin-CreER^T2^; Lgr4^fl/fl^; Vil1-Grem1* animals to assess the post weaning effect of receptor knockout on polyp development and ectopic crypt proliferation. However, receptor knockout was incomplete in these models with faint residual expression of *Fzd5* and *Lgr4* detectable by ISH after recombination (Figure S5A,B). This was felt to be a consequence of incomplete activity of the *Villin-CreER^T2^* recombinase in the undifferentiated progenitor cells of both normal and ectopic crypt bases, and we were able to demonstrate the emergence of unrecombined escaper crypts using *Villin-CreER^T2^; Rosa26^tdTom^; Vil1-Grem1* reporter animals (Figure S5C). Consequently, in *Fzd5* and *Lgr4* knockout models, selection of *Fzd5(+)* and *Lgr4*(+) escaper cells led to repopulation of the intestinal mucosa, restoration of unrecombined levels of tissue expression in developing polyps (Figure S5D) and no long-term impact on overall *Vil1-Grem1* polyp burden or animal survival in either model.

In order to circumvent this problem with *Villin-CreER^T2^* we generated a *Rosa-CreER^T2^*; *Lgr4^fl/fl^; Vil1-Grem1* model, and assessed the impact of acute loss of *Lgr4* receptor expression on ectopic crypt proliferation in 120 day old animals with an established *Vil1-Grem1* phenotype (Figure 5A). Following *Lgr4* receptor knockout, we saw a drop in ectopic crypt cell proliferation 3 days after recombination, implicating environmental wnt ligand-dependency in ectopic crypt maintenance (Figure 5B). Interestingly, there was no equivalent impact on crypt basal cell turnover, implicating a reduced reliance of normal crypts on *Lgr4* receptor signal transduction (Figure 5C). Together, this combination of genetic and pharmacological manipulation of Wnt signalling demonstrates the environmental ligand-dependency of the ectopic niche in early stage lesions, and implicates ectopic crypt epithelial cell expression of Wnt receptors as a key signal transduction component.

**Figure 5.**
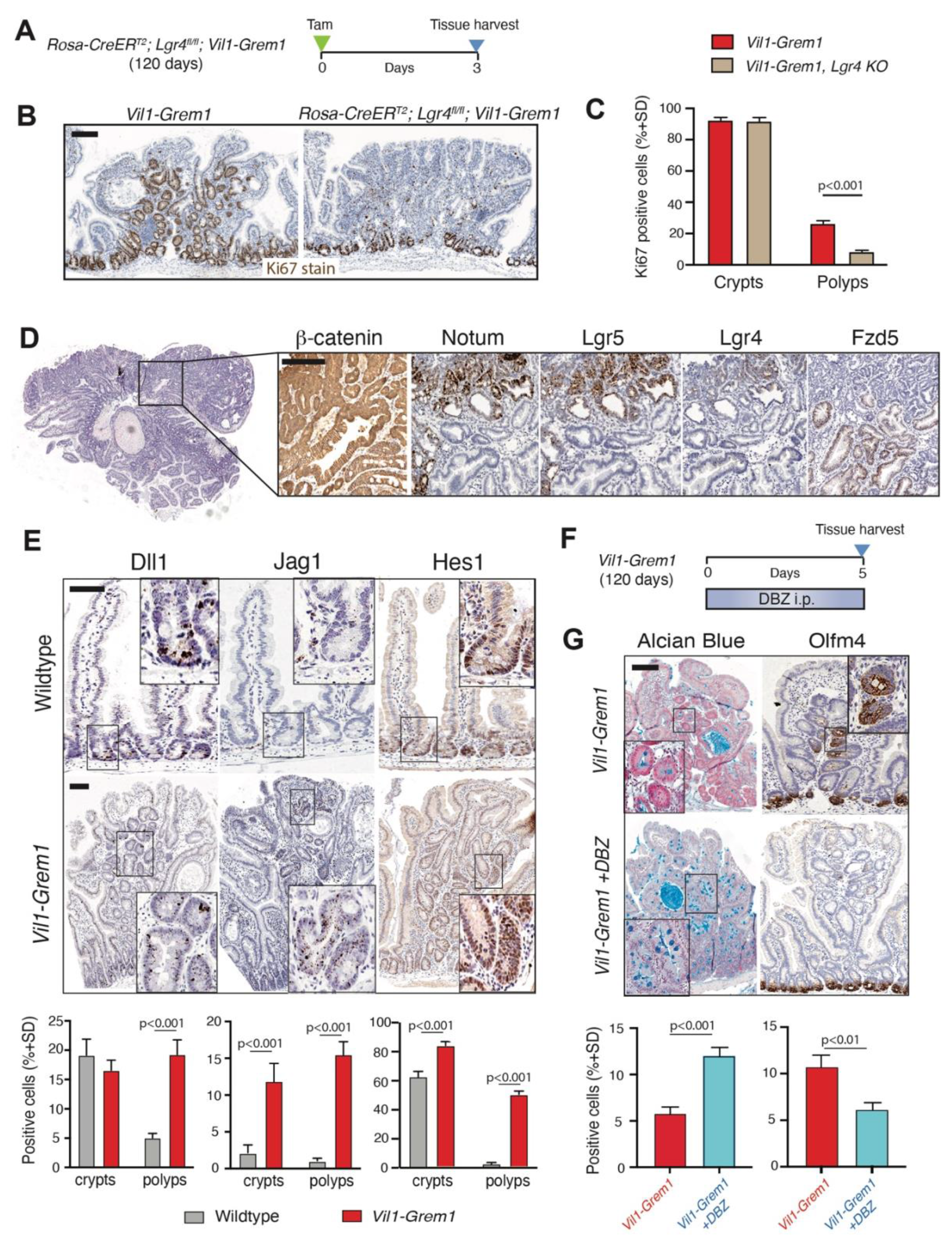
Investigating ectopic niche and advanced polyp signalling. **A.** Schematic shows recombination and harvesting timepoint for acute *Lgr4* knockout in 120 day old *Rosa-CreER^T2^, Lgr4^fl/fl^; Vil1-Grem1* mice. **B.** KI67 staining and **C.** quantification of proliferating crypt base and ectopic crypt cells following acute *Lgr4* knockout (n=5 mice per group, t test, p values as stated). **D.** Change in wnt target and receptor staining in advanced *Vil1-Grem1* polyps. **E.** Representative staining and quantification of Notch pathway ligands (*Dlk1, Jag1*) and target gene (HES1) expression in wildtype and *Vil1-Grem1* mice (n=5 mice per group, t test, p values as stated). **F.** Schematic shows recombination and harvesting timepoint for gamma secretase inhibition with Dibenzazepine in 120- day old *Vil1-Grem1* mice. **G.** Representative staining and quantification of Alcian blue goblet cell staining and OLFM4 expression in untreated and Dibenzazepine-treated *Vil1-Grem1* mice (n=5 mice per group, t test, p values as stated). Scale bars 200µm.

Consistent with previous findings ^12^ we saw membranous b-catenin staining and absence of *Lgr5* expression in the ectopic crypts of early lesions, but did notice nuclear b-catenin and *Lgr5* activation in a small number of large, advanced polyps in older animals. In light of the dependence of ectopic crypt formation on secreted wnt signalling in earlier stage lesions, we hypothesised that these larger polyps somatically acquire wnt disrupting epithelial mutations. Further analysis of these large lesions revealed an overlap of nuclear b-catenin positivity with activation of both *Lgr5* and *Notum* staining and concomitant down-regulation of the wnt receptor *Fzd5* in the regions of these lesions with advanced dysplasia (Figure 5D). Targeted sequencing of dissected advanced polyps revealed epithelial wnt activating mutations in 8/11 (72%) of *Lgr5(+)* lesions with activating b-catenin mutations in seven lesions and a *Ptprk-Rspo3* fusion mutation in one lesion, confirmed with epithelial *RSpo3* staining on ISH (Figure S5E). This frequency of somatic wnt disrupting mutations in large *Vil1-Grem1* polyps with advanced dysplasia is consistent with the previously noted frequency of *APC* mutations in advanced HMPS polyps in humans^12^. Together these data are consistent with a shift away from canonical wnt ligand signal transduction in advanced lesions, and reflects a switch from environmental ligand-dependency to acquired epithelial cell wnt autonomy as polyps progress. Notably, these advanced lesions also downregulated epithelial *Grem1* expression, through loss of the *Villin* differentiation marker in high-grade dysplasia, and acquired constitutive wnt activation through epithelial mutation rendering them insensitive to both UCB Ab7326 antibody and porcupine effect.

Together these data demonstrate reversible dependence of ectopic crypt stem cell function and proliferation on secreted, microenvironmental wnt ligand activity. Remodelled stroma acts as a key source of wnt ligand expression, through *Grem1*-induced de-repression of stem cell niche supporting fibroblasts. Although this can be therapeutically blocked by porcupine inhibition which successfully prevents villus stem cell activity, inhibition of wnt in the ectopic niche does not lead to comparable resolution of the stromal functional architecture induced through inhibition of paracrine *Grem1*.

### Ectopic crypt Notch pathway activation helps maintain aberrant stem cell populations

Given that tissue staining had identified aberrant OLFM4 expression specifically within ectopic crypts in untreated *Vil-1-Grem1* animals, we next explored the Notch pathway, as OLFM4 is a direct Notch target gene. ISH staining revealed elevated expression of canonical (*Jag1*) and non-canonical ligands (*Dlk1*), with upregulation of the Notch target gene HES1 (IHC) in the ectopic crypt cells indicating Notch pathway activity (Figure 5E). Short term inhibition (5 days) of the pathway with the 𝛄-secretase inhibitor Dibenzazepine (DBZ) ^17^, had no macroscopic impact on the number of ectopic crypts or villus cell proliferation (KI67 stain) (Figure 5F). However, it did result in loss of ectopic crypt OLFM4 staining and significant increase in the number of Alcian blue stained goblet cells in ectopic crypts (Figure 5G). Notably, porcupine inhibition of wnt activity also led to a significant reduction in expression of *Jag1* and OLFM4 expression in the villus compartment (Figure S5F). This indicated a role for Notch in Lgr5(-) Olfm4(+) stem cell maintenance and binary fate selection in the ectopic crypt, and indicated that this activity was downstream of secreted wnt activation.

### Human polyps

Having used a disease-positioned mouse model to identify the importance of stromal remodelling in the generation of wnt dependent ectopic stem cell niche, we turned to equivalent human samples to find evidence of the same processes. We and others have previously shown that HMPS polyps and a proportion of sporadic TSA lesions have marked ectopic crypt formation with aberrant epithelial expression of Grem1^12^. These lesions progress to an advanced polyp stage through acquired wnt disruption, including a high frequency of ligand-dependent *R-Spondin* fusions and *Rnf43* mutations in TSA’s^18^. In contrast, conventional tubulovillous adenomas arise through constitutive epithelial wnt activation as a consequence of initiating *APC* mutation in crypt base columnar cells, and have only stromal expression of *Grem1* (Figure S6). In light of the preclinical findings identifying a role for stromal remodelling and wnt ligand dependency we assessed some very small human HMPS, TSA and conventional adenoma lesions to look for evidence of de-repressed *CD55(+) WNT2b(+)* fibroblast cells. We were able to identify these cells in significant numbers surrounding ectopic crypts in HMPS and TSA lesions, but could find very few within the stroma of the conventional TVA’s (Figure 6A,B). This indicates that stromal remodelling with de-repression of a *CD55(+) WNT2b(+)* stromal cell population is a shared feature in mouse and human lesions characterised by ectopic crypt formation, but is not seen in Wnt ligand-independent tubulovillous adenomas where constitutive epithelial wnt activation negates the need for environmental wnt ligand supply.

**Figure 6.**
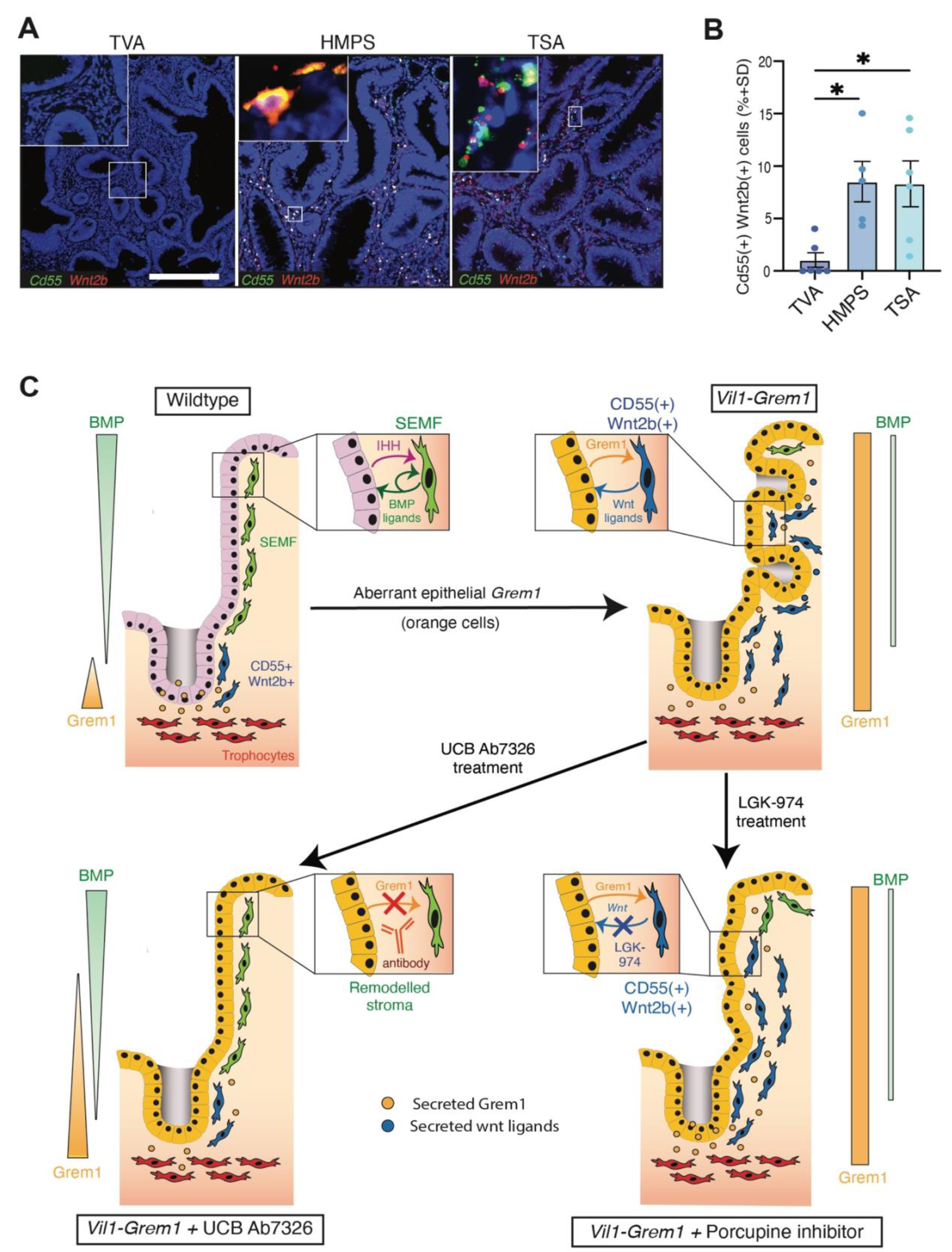
Human polyp stromal remodelling and model summary. **A.** Representative fluorescent co- ISH staining images, and **B.** quantification of *CD55(+) WNT2B(+)* cell proportions in the stroma of small sporadic tubulovillous adenoma (TVA n=6), Hereditary Mixed Polyposis Syndrome (HMPS n=5) and sporadic traditional serrated adenoma (TSA n=6) lesions. **C.** Model summary depicting the disruption of BMP signalling gradients leading to a remodelling of the stromal microenvironment and establishment of wnt ligand-dependent ectopic crypt stem cell niches. Scale bar 100µm.

## Discussion

Strictly controlled, counterbalanced secreted signalling pathways underpin the unique structure and function of the intestinal mucosa. Continual epithelial-mesenchymal crosstalk defines mucosal architecture during organ development^7^ and underpins tissue homeostasis. The impact of this interaction on homeostatic epithelial cell fate is long established ^3^, however, the role of morphogens in determining stromal cell function is less well understood. Recently, the Shivdasani group have explored the anatomy of intestinal stromal cell heterogeneity, demonstrating a functionally layered fibroblast structure^8^, dependent on cell position within mesenchymal-secreted BMP signalling gradients. In this model, proximity to a luminal surface BMP ligand source suppressed the stromal expression of stem cell niche supporting genes, with appropriate de-repression only occurring at the crypt base through distance from the luminal BMP ligand source and the sub-crypt presence of secreted BMPi ^8^. This has been further supported by the development of a novel model system to mechanistically interrogate the study of epithelial-fibroblast self-organisation *in vitro* ^19^. Lin et al demonstrate the requirement for stromal BMP responsiveness for the subsequent development of epithelial crypt-like structures, indicating a key role for stromal cell organisation in promoting epithelial cell fate polarisation and crypt base niche formation^19^. The sensitivity of the stromal cells to BMP regulation was further demonstrated in vivo by Ouahoud et al who used deletion of stromal *Bmpr1a* receptors to alter the stromal cell secretome and initiate polyp formation^20^. Together, this work represents a shift in the epithelial-centric intestinal cell fate paradigm to a co-evolutionary model, where the interface between functionally distinct epithelial and stromal cell populations is established through shared exposure to paracrine signalling concentration gradients. Thus, crypt-villus compartmentalisation is dependent on bi-directional crosstalk between epithelial and stromal cells, with each cell population influencing and regulating the fate and function of each other. This cell compartment interdependence enables the tissue to be exquisitely sensitive and responsive to local change, which is required for a rapid and adaptive physiological response to damage^21^, but also renders the mucosa susceptible to pathological dysregulation, especially following mutation-induced disruption of the key regulatory pathways.

Here, we used a disease-positioned model of a rare human polyposis syndrome to mechanistically explore the impact of pathological disruption of polarised BMP gradients and provide insight into the co-dependence of epithelial-mesenchymal fate and function in the gut. Using a series of specific pathway inhibitors, we were able to triangulate the spatio-temporal consequences of cumulative signalling pathway disruption and define a pathological sequence of events. We demonstrate that aberrant pan-epithelial *Grem1* expression acts intercompartmentally to disrupt BMP signalling gradients in both epithelium and underlying stromal cell populations. Resultant remodelling of the stromal cell compartment results in the de-repression and expansion of *Cd55(+) Wnt2b(+)* fibroblasts capable of supporting stem cell function. Ultimately this leads to the formation of a wnt ligand-dependent ectopic stem cell niche outside of the confines of the crypt base. Wnt-stimulated epithelial Notch activity within the ectopic crypts then co-supports SOX9(+) OLFM4(+) Lgr5(-) stem cell function. Although abrogation of wnt ligands through porcupine inhibition does prevent ectopic crypt formation and significantly suppresses advanced polyp formation, it is only attenuation of aberrant paracrine BMP antagonism that is capable of reverting fibroblast remodelling and allowing re-imposition of appropriate homeostatic stromal cell functional segregation. The dependence of this permissive ectopic stem cell niche on paracrine signalling renders the phenotype therapeutically reversible in advance of acquisition of epithelial somatic mutation. Together this work demonstrates that aberrant epithelial *Grem1* expression disrupts homeostatic BMP crosstalk between the epithelium and the underlying mesenchyme, and transforms the mucosal stromal cell architecture into a microenvironment capable of sustaining an ectopic stem/progenitor cell population. This illustrates the pathological consequence of disruption of homeostatic morphogen gradients in a human disease relevant setting, and supports an emerging co-evolutionary model of intestinal cell compartmentalisation based on bidirectional regulation of epithelial-mesenchymal cell fate and function (Figure 6C).

In established cancers, *GREM1* expression is frequently upregulated in the desmoplastic stroma of a number of solid tumours ^22^. In sporadic colorectal cancer, high stromal *GREM1* expression is clearly associated with aggressive mesenchymal subtypes and poor prognosis ^12^. As we have demonstrated a role for *GREM1* in remodelling the stromal microenvironment to generate an ectopic stem cell niche in a disease-positioned model of a hereditary intestinal polyposis, it would be interesting to explore whether this mechanism persists into established sporadic cancers. This represents a potential therapeutic opportunity given the increasing interest in the role of the tumour microenvironment in supporting both *LGR5*(+) and (-) cancer stem cell populations ^23,24^. Other recent publications have also implicated a role for *Grem1* in activation of fibroblast growth factor receptor signalling^25^ and in regulating epithelial mesenchymal transition^26^ in prostate and pancreatic cancer respectively. Together this provides accumulating evidence for a possible pleiotrophic role for *GREM1* inhibition in neoplasia and the UCB anti-Grem1 antibody has entered into human clinical trials.

What is clear from this work is that the remarkable phenotypic reversion and long-term survival of Vil1-Grem1 mice commenced early on UCB Ab7326 antibody therapy, provides hope that *Grem1* inhibition could offer a preventative therapeutic strategy for HMPS patients. This hereditary polyposis carries a high lifelong cancer risk and prophylactic treatment could potentially reduce the need for regular intrusive surveillance colonoscopies or prophylactic colectomy in this rare patient population.

### Grant support

SJL was supported by CRUK Programme Grant (DRCNPG-Jun22\100002) and a Wellcome Trust (Senior Clinical Research Fellowship (206314/Z/17/Z). EJM was supported by the Lee Placito Research Fellowship (University of Oxford). JEE was supported by National Institute for Health Research (NIHR) Oxford Biomedical Research Centre. Experimental work was also supported by Rosetrees Trust and Stoneygate Trust research grant (M493) and a UCB Pharma research grant. Animal costs were supported by an International Accelerator Award, ACRCelerate, jointly funded by Cancer Research UK (A26825 and A28223), FC AECC (GEACC18004TAB) and AIRC (22795), and the MRC Mouse Genetics Network. Core funding to the Wellcome Centre for Human Genetics was provided by the Wellcome Trust (090532/Z/09/Z).). OJS, NG, MH and RR were supported by CRUK core funding to the CRUK Scotland Institute (A31287) and core programme awards to OJS (A21139 and DRCQQR- May21\100002). This research was funded in part, by the Wellcome Trust. For the purpose of Open Access, the author has applied a CC BY public copyright licence to any Author Accepted Manuscript version arising from this submission.

## Author Contributions

EJM, HBD, IT and SJL conceived and designed the project. Funding obtained by SJL and EJM. Experiments were conducted by HBD, EJM, NG, AL, SB, EG, SO, NN, RR, and MH. Bioinformatic analysis carried out by GV. Pathology support LMW. Tissue, materials, and data provision KSM, LKJ, JEE, LMW, ND, TP, GD. Conceptual input and data interpretation GD, ND, OS, IT. Manuscript written by EJM and SJL.

## Disclaimer

The views expressed are those of the author/s and not necessarily those of the NHS, the NIHR or the Department of Health

## Acknowledgements

The authors thank patients visiting the John Radcliffe Hospital, especially those with HMPS, for their generous tissue donation through the Oxford TGU GI Illness Biobank. The authors also thank Derek Magee from HeteroGenius for digital reconstruction of lineage tracing ribbons in polyps. The authors are grateful to the Core Research Services at the CRUK Scotland Institute: Biological Services Unit and Histology services for technical support and to Catherine Winchester for critical reading of the manuscript.

## Materials and Methods Resource availability

### Lead contacts

Further information and requests for resources and reagents should be directed to and will be fulfilled by the lead contacts, Eoghan J Mulholland Hayley Belnoue-Davis or Simon J Leedham.

### Materials availability

Mouse lines generated in this study are available with an MTA.

### Data availability

Raw image/micrograph files which support the findings in this study are available from the corresponding authors upon request.

### Experimental model and subject details Animals

All procedures were carried out in accordance with Home Office UK regulations and the Animals (Scientific Procedures) Act 1986 as described previously^23^. All mice were housed in individually ventilated cages at the animal unit either at the Functional Genetics Facility (Wellcome Centre for Human Genetics, University of Oxford) or The CRUK Scotland Institute (Glasgow). All mice were housed in a specific-pathogen-free (SPF) facility, with unrestricted access to food and water, and were not involved in any previous procedures. All strains used in this study were maintained on C57BL/6J background for ≥6 generations. All procedures were carried out on mice of at least 6 weeks of age, both male and female.

### Generation of *Rosa26^Grem1^* Mouse

To generate *Rosa26^Grem1^* mice, a Grem1 complementary DNA cassette was cloned into the integrase mediated cassette exchange vector (CB93) and used to transfected into RS-PhiC31 ES cells. Recombinant clones were obtained which harbored the Grem1 cDNA transgene positioned within the Rosa26 locus, allowing for Cre-dependent activation of transgene expression. Recombinant clones were injected into blastocysts, and chimeras were generated (Wellcome Centre for Human Genetics Trangenics Core). Chimeras were crossed with wild-type C57BL/6J mice to obtain F1 heterozygotes.

### Human subjects

FFPE sections of Human TSA, TVA and HMPS lesions were collected from the Oxford GI biobank OCHRe: 22/A158 and 22/A158b. All samples were subject to expert histopathological review.

### Methods Details Treatment of animals

The mouse alleles used in this study in various combinations are as follows: *Vil1-Grem1*^12^*, Atoh1CreER^T2^* ^13^*, Villin-CreER^T2^* ^27^*, RosaCreER^T2^* ^28^*, Sox9CreER^T2^* ^29^ *Rosa26^Grem1^, Rosa26^tdTom^* ^30^ *Rosa26^YFP^* ^31^*, Lgr4^fl/fl^*^32^*, Fzd5^fl/fl^* ^33^. All experimental mice were from a C57/BL6 background, backcrossed for at least 6 generations, and were housed in specific-pathogen-free cages. For inducible models, recombination was achieved using the free base tamoxifen (Sigma-Aldrich, St. Louis, MO) dissolved in ethanol/oil (1:9). For lineage tracing experiments (*Atoh1CreER^T2^;Rosa26^tdTom^* and *Atoh1CreER^T2^;Rosa26^Grem1^*), recombination was induced by a single dose of 3 mg of tamoxifen. For anti-Grem1 therapy mice received weekly 30mg/kg subcutaneous administration of UCB Ab7326 (UCB Pharma). To generate Kaplan-Meier data, mice were sacrificed when they reached humane-end points (exhibited anaemia, hunching and inactivity). For Wnt blocking LGK-974 (MedKoo, 205851) was used. LGK-974 was administered at a concentration of 5 mg/kg, in 0.5% Methylcellulose twice daily by oral gavage. For Notch signalling inhibition experiments, DBZ (Gamma-secretase inhibitor) was used at a dose of 30 mmol/kg.

### Formalin-fixed paraffin embedded processing

Gut preparations were washed in PBS, fixed overnight in 10% neutral buffered formalin and then transferred to 70% ethanol before processing for embedding. Formalin-fixed gut sections were rolled into Swiss Rolls, pinned and placed in a histology cassette. Specimens were processed using a Histomaster machine (Bavimed). Processed samples were embedded in paraffin wax using a paraffin embedding station (EG1150H, Leica)^23^.

### In situ hybridisation

For in situ hybridisation (ISH) of both human and mouse FFPE samples, 4 μm formalin-fixed, paraffin-embedded tissue sections were used as previously described^23^. The sections were baked at 60°C for 1 h before dewaxing in xylene and ethanol. Chromogenic ISH was performed using the RNAscope^®^ 2.5 HD -BROWN and Fluorescent ISH was then performed using the RNAscope Fluorescent Multiplex Reagent Kit (Bio-techne) in accordance with the supplier’s guidelines. All probes were purchased from ACD (Bio-techne),and included: Mm-Wnt5a (ACD, Cat# 316791); Mm-Rspo2 (ACD, Cat# 402001); Mm- Wnt2b (ACD, Cat# 405031); Mm-Notum (ACD, Cat# 428981); Mm-Lgr4 (ACD, Cat# 318321); Mm-Fzd5 (ACD, Cat# 404911); Mm-Dll1 (ACD, Cat# 425071); Mm-Jag1 (ACD, Cat# 412831); Mm-Axin2 (ACD, Cat# 400331); Hs-CD55 (ACD, Cat# 426551); Hs-Wnt2b (ACD, Cat# 453361-C2).

### Immunohistochemistry

Sections were de-paraffinised in xylene and rehydrated through graded alcohols to water. Antigen retrieval was achieved by pressure cooking in 10 mmol/L citrate buffer (pH 6.0) for 5 min. Endogenous peroxidase activity was blocked by incubating in 3% hydrogen peroxidase (in methanol) for 20 min. Next, sections were blocked with 1.5% serum for 30 min, after which they were incubated with primary antibodies for 1 h. The sections were then incubated with appropriate secondary antibodies for 30 min at room temperature. For chromogenic visualisation, sections were incubated with ABC (Vector labs) for 30 min and stained using DAB solution (VectorLabs), after which they were counterstained with hematoxylin, dehydrated and mounted^23^. All antibodies used in this work are as follows: Anti-Ki67 (D3B5) Rabbit mAb (Cell Signalling Technology, Cat#CS12202S); Anti-lysozyme Rabbit pAb (DAKO, Cat#EC3.2.1.17); Anti-mCherry (TdTomato) mouse mAb (Novus Bio, Cat#NBP1- 96752); Anti-SOX9 Rabbit pAb (Sigma Aldrich, Cat#AB5535); Human/Mouse EphB2 Antibody (R&D, Cat#AF467; Phospho-SMAD1/5 (Ser463/465) Rabbit mAb (Cell Signalling Technology, Cat#41D10); Anti-β-Catenin Clone 14 (BD Biosciences, Cat#610154); HES1 (D6P2U) Rabbit mAb (Cell Signalling Technology, Cat#11988); Olfm4 (D6Y5A) XP® Rabbit mAb (Cell Signalling Technology, Cat#39141); anti-RFP Rabbit pAb (Rockland, Cat#600-401-379).

### Multiplex immunofluorescence

Multiplex immunofluorescence (MPIF) staining was performed on FFPE sections of thickness 4-μm using the OPAL protocol (Akoya Biosciences, Marlborough, MA) on the Leica BOND RXm autostainer (Leica Microsystems, Wetzlar, Germany)^23^. Six consecutive staining cycles were performed using the following primary antibody-Opal fluorophore pairs:

### Immune panel

1. Ly6G (1:300, 551459; BD Pharmingen)–Opal 540; (2) CD4 (1:500, ab183685;

Abcam)–Opal 520; (3) CD8 (1:800, 98941; Cell Signaling)–Opal 570; (4) CD68 (1:1200, ab125212; Abcam)–Opal 620; (5) FoxP3 (1:400, 126553; Cell Signaling)–Opal 650; and (6) E-cadherin (1:500, 3195; Cell Signaling)–Opal 690.

### Matrix panel

1. Laminin (1:400, ab11575; Abcam)-Opal 540; (2) Tenascin-C(1:600, ab108930; Abcam)-Opal 520; (3) Fibronectin (1:1000, F3648; Sigma-Aldrich)-Opal 570; (4) Osteopontin (1:750, ab218237; Abcam)-Opal 620; MMP3 (1:100,ab52915; Abcam)-Opal 650; (5) Collagen I (1:400, 72026; Cell Signaling)-Opal 690.

Tissues sections were incubated for 1 h in primary antibodies and detected using the BOND Polymer Refine Detection System (DS9800; Leica Biosystems, Buffalo Grove, IL) in accordance with the manufacturer’s instructions, substituting DAB for the Opal fluorophores, with a 10-min incubation time and withholding the hematoxylin step. Antigen retrieval at 100°C for 20 min, in accordance with standard Leica protocol, with Epitope Retrieval Solution one or two was performed prior to each primary antibody being applied. Sections were then incubated for 10 min with spectral DAPI (FP1490, Akoya Biosciences) and the slides mounted with VECTASHIELD Vibrance Antifade Mounting Medium (H-1700-10; Vector Laboratories). Whole-slide scans and multispectral images (MSI) were obtained on the Akoya Biosciences Vectra Polaris. Batch analysis of the MSIs from each case was performed with the inForm 2.4.8 software provided. Finally, batched analysed MSIs were fused in HALO (Indica Labs) to produce a spectrally unmixed reconstructed whole-tissue image. Cell density analysis was subsequently performed for each cell phenotype across the three MPIF panels using HALO.

### HiPlex ISH

HiPlex ISH staining was performed on FFPE sections of thickness 4-μm outsourced to Bio-Techne who used the RNAscope™ HiPlex12 Reagent Kit (488, 550, 650, 750) v2 Standard Assay. This panel probed for the following mouse RNA transcripts: Mm-Wnt2b-T6 (ACD, Cat#405031-T6); Mm-Bmp5-T7 (ACD, Cat#401241-T7); Mm-Cd81-T5 (ACD, Cat#556971-T5); Mm-Postn-T8 (ACD, Cat#418581-T8); Mm- Rspo3-T2 (ACD, Cat#402011-T2); Mm-Grem1-T3 (ACD, Cat#314741-T3); Mm-Serpine1-T1 (ACD, Cat#402501-T1); Mm-Cd55-T4 (ACD, Cat#421251-T4); Mm-Pdgfra-T10 (ACD, Cat#480661-T10); Mm- Cd3e-T11 (ACD, Cat#314721-T11); Mm-Acta2-T9 (ACD, Cat#319531-T9); Mm-Cdh1-T12 (ACD, Cat#408651-T12).Whole-slide scans and multispectral images (MSI) were obtained on the Akoya Biosciences Vectra Polaris.

### Image analysis

IHC images were analysed as follows: Positive cells were quantified using QuPath digital pathology software (v0.2.3, (Bankhead et al., 2017)), downloaded from https://QuPath.github.io/. Firstly, annotations of tissue areas were created for each sample with areas of folded tissue excluded to eliminate false positive signals. Cells were identified within QuPath using a custom algorithm established via stain separation using color reconstruction. Positive cell detection analysis was run to identify DAB positive cells and results reported as percentage of positive cells. Each annotation was manually verified for correct signal identification. For analysis of multiplex IHC, HiPlex ISH and dual ISH, HALO image analysis software (Indica Labs) was used to identify cell phenotypes, cell density analysis and, mapping of cell phenotypes onto images. For 3D reconstruction of linage traced crypts in Vil1-Grem1-Sox9YFP polyps, images were aligned using HeteroGenius MIM with 3D Pathology AddOn (HeteroGenius, Leeds, UK).

### Statistical analysis

All statistical calculations were performed using GraphPad Prism 10. Unpaired two-tailed t tests were used to determine statistical significance between two groups. One-way or two-way analysis of variance (ANOVA) was used to assess statistical significance between three or more groups. For precise details on statistical analyses (including post-hoc tests), size of n, tests utilised, and significance definition (p-values) see corresponding figure legend and results.

## References

1 Scoville, D., Sato, T., He, X. & Li, L. Current view: intestinal stem cells and signaling. Gastroenterology 134, 849–864 (2008).

2 van den Brink, G. & Offerhaus, G. The morphogenetic code and colon cancer development. Cancer Cell 11, 109–117 (2007).

3 McCarthy, N., Kraiczy, J. & Shivdasani, R. A. Cellular and molecular architecture of the intestinal stem cell niche. Nat Cell Biol 22, 1033–1041 (2020). 10.1038/s41556-020-0567-z

4 Biswas, S. et al. Microenvironmental control of stem cell fate in intestinal homeostasis and disease. J Pathol (2015). 10.1002/path.4563

5 McCarthy, N. et al. Distinct Mesenchymal Cell Populations Generate the Essential Intestinal BMP Signaling Gradient. Cell Stem Cell 26, 391–402.e395 (2020). 10.1016/j.stem.2020.01.008

6 McCarthy, N. et al. Smooth muscle contributes to the development and function of a layered intestinal stem cell niche. Dev Cell 58, 550–564.e556 (2023). 10.1016/j.devcel.2023.02.012

7 Shyer, Amy E., Huycke, Tyler R., Lee, C., Mahadevan, L. & Tabin, Clifford J. Bending Gradients: How the Intestinal Stem Cell Gets Its Home. Cell 161, 569–580 (2015). 10.1016/j.cell.2015.03.041

8 Kraiczy, J. et al. Graded BMP signaling within intestinal crypt architecture directs self-organization of the Wnt-secreting stem cell niche. Cell Stem Cell 30, 433–449.e438 (2023). 10.1016/j.stem.2023.03.004

9 Jaeger, E. et al. Hereditary mixed polyposis syndrome is caused by a 40-kb upstream duplication that leads to increased and ectopic expression of the BMP antagonist GREM1. Nat Genet (2012). ng.2263 [pii] 10.1038/ng.2263

10 Haramis, A. P. et al. De novo crypt formation and juvenile polyposis on BMP inhibition in mouse intestine. Science 303, 1684–1686 (2004).

11 Batts, L. E., Polk, D. B., Dubois, R. N. & Kulessa, H. Bmp signaling is required for intestinal growth and morphogenesis. Dev Dyn 235, 1563–1570 (2006). 10.1002/dvdy.20741

12 Davis, H. et al. Aberrant epithelial GREM1 expression initiates colonic tumorigenesis from cells outside the stem cell niche. Nat Med 21, 62–70 (2015). nm.3750 [pii] 10.1038/nm.3750

13 Tomic, G. et al. Phospho-regulation of ATOH1 Is Required for Plasticity of Secretory Progenitors and Tissue Regeneration. Cell Stem Cell 23, 436–443.e437 (2018). 10.1016/j.stem.2018.07.002

14 Jaeger, E. et al. Common genetic variants at the CRAC1 (HMPS) locus on chromosome 15q13.3 influence colorectal cancer risk. Nat Genet 40, 26–28 (2008).

15 Davies, G. C. G. et al. Discovery of ginisortamab, a potent and novel anti-gremlin-1 antibody in clinical development for the treatment of cancer. MAbs 15, 2289681 (2023). 10.1080/19420862.2023.2289681

16 Kapoor, V. N. et al. Gremlin 1(+) fibroblastic niche maintains dendritic cell homeostasis in lymphoid tissues. Nat Immunol 22, 571–585 (2021). 10.1038/s41590-021-00920-6

17 van Es, J. H., de Geest, N., van de Born, M., Clevers, H. & Hassan, B. A. Intestinal stem cells lacking the Math1 tumour suppressor are refractory to Notch inhibitors. Nature Communications 1, 18 (2010). 10.1038/ncomms1017

18. Sekine, S., Yamashita, S., Tanabe, T., Hashimoto, T. & Yoshida, H. Frequent PTPRK-RSPO3 fusions and RNF43 mutations in colorectal traditional serrated adenoma. Jornal of Pathology In Press (2016).

19 Lin, M. et al. Establishment of gastrointestinal assembloids to study the interplay between epithelial crypts and their mesenchymal niche. Nat Commun 14, 3025 (2023). 10.1038/s41467-023-38780-3

20 Ouahoud, S., Hardwick, J. C. H. & Hawinkels, L. Extracellular BMP Antagonists, Multifaceted Orchestrators in the Tumor and Its Microenvironment. Int J Mol Sci 21 (2020). 10.3390/ijms21113888

21 Koppens, M. A. J. et al. Bone Morphogenetic Protein Pathway Antagonism by Grem1 Regulates Epithelial Cell Fate in Intestinal Regeneration. Gastroenterology 161, 239–254.e239 (2021). 10.1053/j.gastro.2021.03.052

22 Sneddon, J. et al. Bone morphogenetic protein antagonist gremlin 1 is widely expressed by cancer-associated stromal cells and can promote tumor cell proliferation. Proc Natl Acad Sci U S A 103, 14842–14847 (2006).

23 Vasquez, E. G. et al. Dynamic and adaptive cancer stem cell population admixture in colorectal neoplasia. Cell Stem Cell 29, 1213–1228.e1218 (2022). 10.1016/j.stem.2022.07.008

24 Cañellas-Socias, A. et al. Metastatic recurrence in colorectal cancer arises from residual EMP1(+) cells. Nature 611, 603–613 (2022). 10.1038/s41586-022-05402-9

25 Cheng, C. et al. Gremlin1 is a therapeutically targetable FGFR1 ligand that regulates lineage plasticity and castration resistance in prostate cancer. Nat Cancer 3, 565–580 (2022). 10.1038/s43018-022-00380-3

26 Lan, L. et al. GREM1 is required to maintain cellular heterogeneity in pancreatic cancer. Nature 607, 163–168 (2022). 10.1038/s41586-022-04888-7

27 el Marjou, F., et al. Tissue-specific and inducible Cre-mediated recombination in the gut epithelium. Genesis 39, 186–193 (2004).

28 Hameyer, D. et al. Toxicity of ligand-dependent Cre recombinases and generation of a conditional Cre deleter mouse allowing mosaic recombination in peripheral tissues. Physiol Genomics 31, 32–41 (2007). 10.1152/physiolgenomics.00019.2007

29 Akiyama, H. et al. Osteo-chondroprogenitor cells are derived from Sox9 expressing precursors. Proc Natl Acad Sci U S A 102, 14665–14670 (2005). 10.1073/pnas.0504750102

30 Madisen, L. et al. A robust and high-throughput Cre reporting and characterization system for the whole mouse brain. Nat Neurosci 13, 133–140 (2010). 10.1038/nn.2467

31 Srinivas, S. et al. Cre reporter strains produced by targeted insertion of EYFP and ECFP into the ROSA26 locus. BMC Developmental Biology 1, 4 (2001). 10.1186/1471-213X-1-4

32 Kato, S. et al. Leucine-rich repeat-containing G protein-coupled receptor-4 (LGR4, Gpr48) is essential for renal development in mice. Nephron Exp Nephrol 104, e63-75 (2006). 10.1159/000093999

33 van Es, J. H. et al. Wnt signalling induces maturation of Paneth cells in intestinal crypts. Nature Cell Biology 7, 381–386 (2005). 10.1038/ncb1240

