## Supplementary Figures for "GREMLIN1 disrupts intestinal epithelial-mesenchymal crosstalk to induce a wnt-dependent ectopic stem cell niche via stromal remodelling"

**Supplementary Figures and Figure Legends**

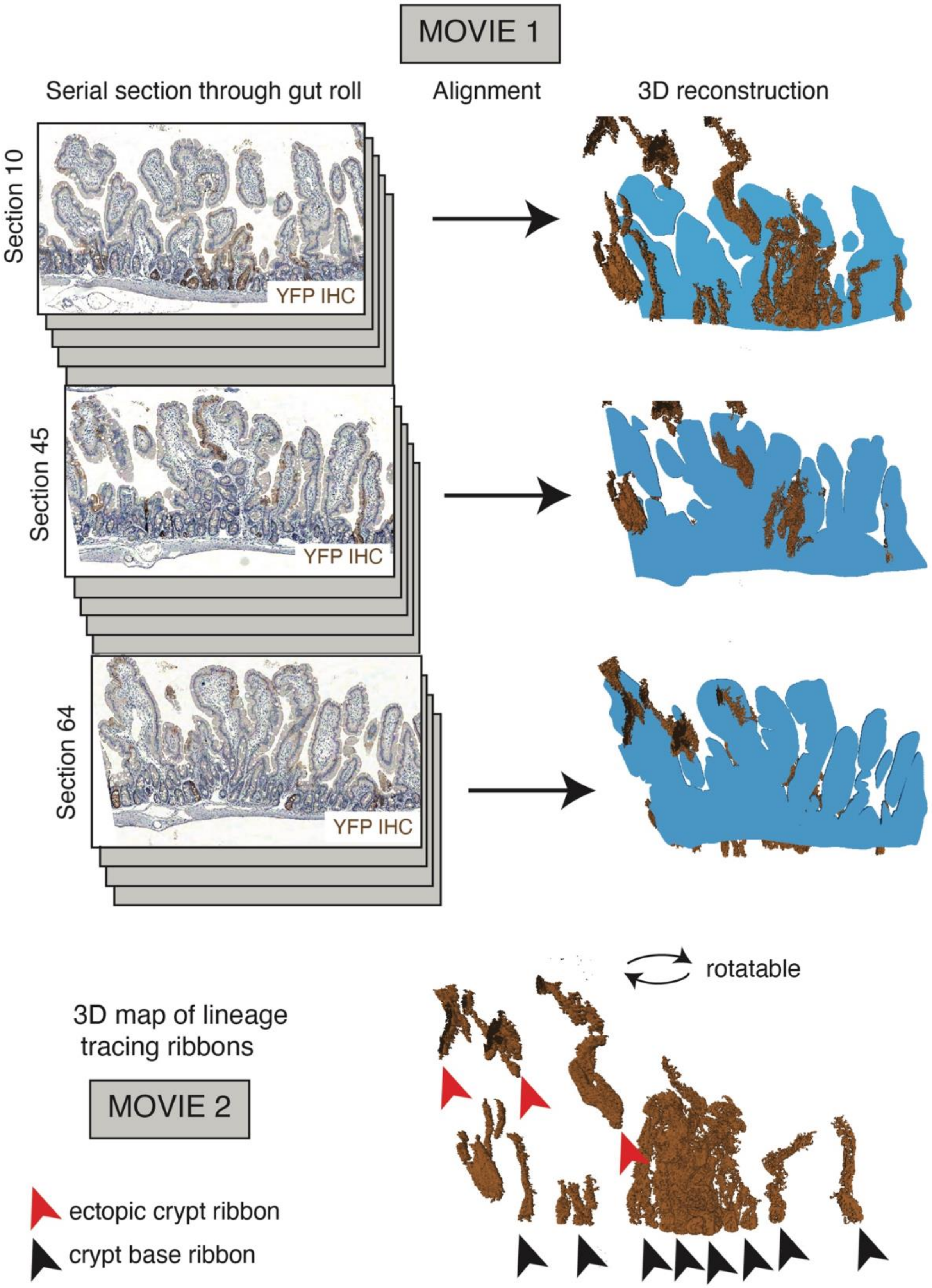

**Supplementary Figure 1. 3D reconstruction to track lineage tracing ribbons through polyps.** Careful serial sectioning through gut rolls with automated anti-YFP staining (brown). Tissue alignment (HeteroGenius, Leeds, UK) and reconstruction of polyps allowed 3-dimensional tracking of lineage tracing ribbons (brown) to show that ectopic crypt ribbons (red arrowheads) were spatially distinct from those arising from the crypt base (black arrowheads).

**Supplementary Movie 1.** Alignment and scrolling of serial sections through >120 day old *Sox9-CreER*$^{T2}$*; Rosa26*$^{YFP}$*; Vil1-Grem1* mouse small intestinal gut roll with anti YFP staining (brown) and automated false colourisation of tracing ribbons (green).

**Supplementary Movie 2.** Rotation of reconstructed polyp to show spatial segregation of crypt basal and ectopic crypt lineage tracing ribbons, to exclude interconnection of ribbons in three-dimensional space.

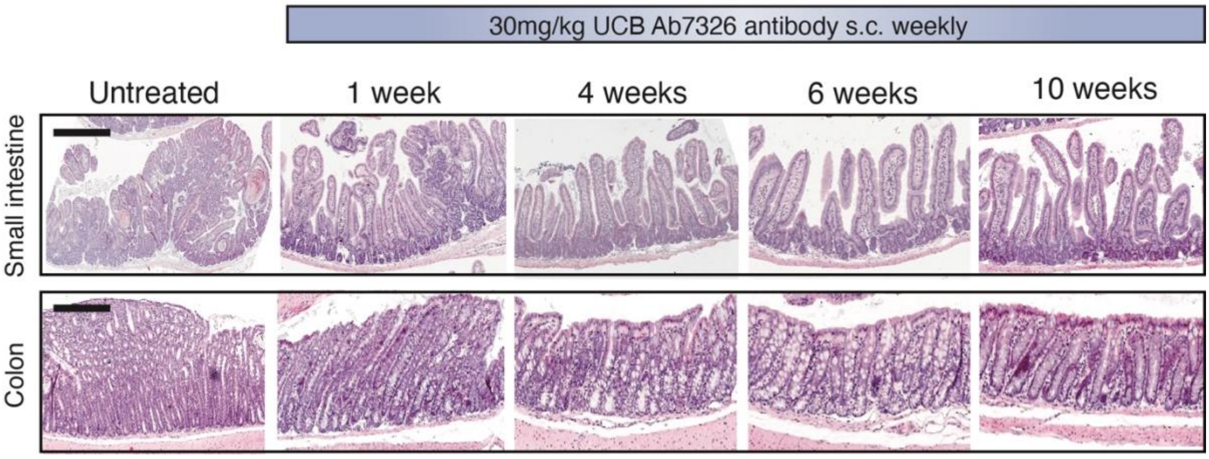


**Supplementary Figure 2. UCB Ab7326 treatment reverses *Vil1-Grem1* mouse polyposis phenotype.** Representative H&E stains of proximal small intestine and colon of *Vil1-Grem1* animals following variable time treatment with UCB Ab7326 antibody. Scale bars 200µm.

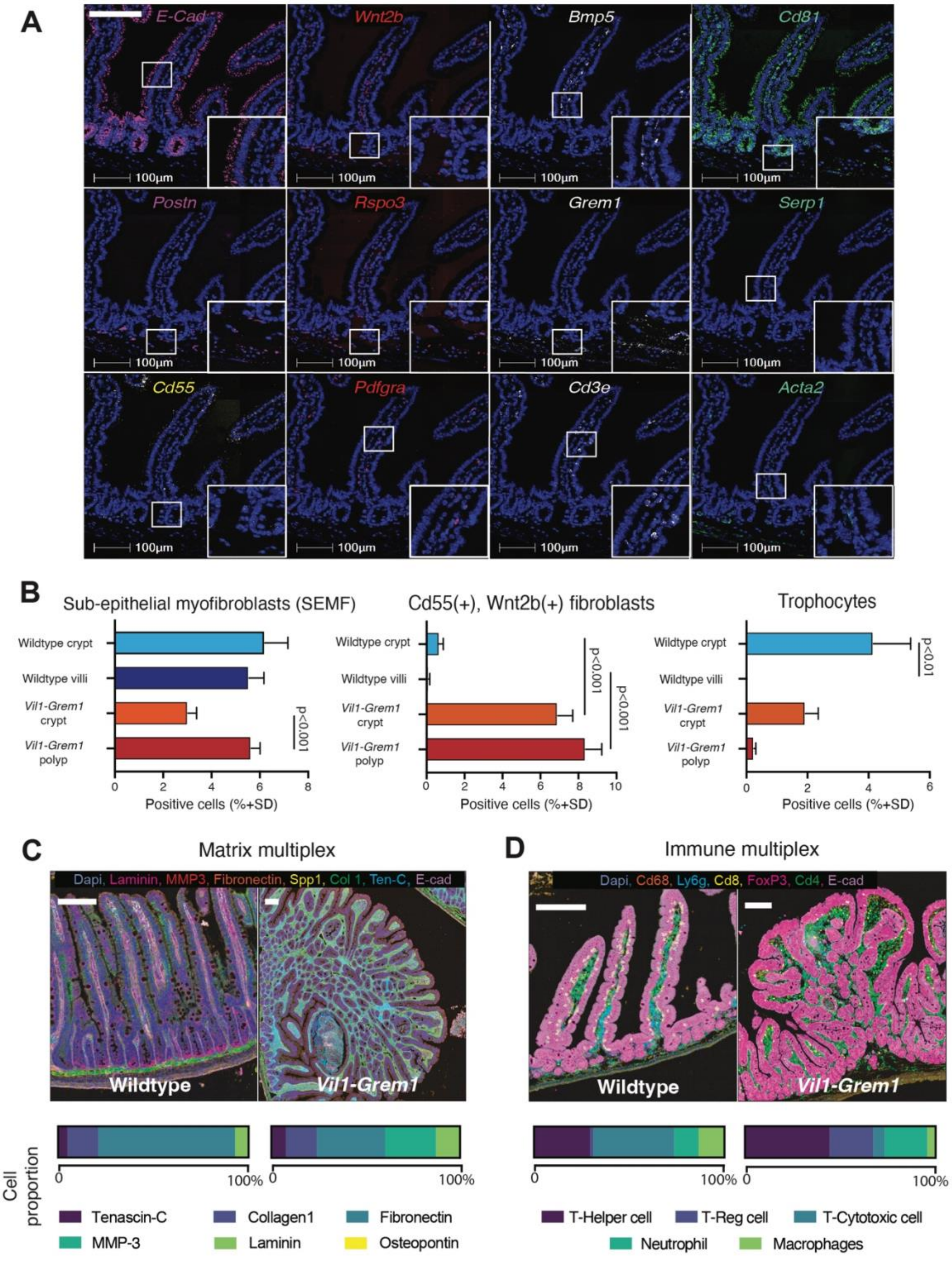


**Supplementary Figure 3. Multiplex ISH and IHC imaging of *Vil1-Grem1* mouse phenotype. A.** Intestinal fibroblast custom Hi-plex panel - individual ISH stain of panel constituents. **B**. Quantification of Hi-plex staining to identify different fibroblast phenotypes in crypts and villi/polyps in wildtype or established *Vil1-Grem1* animals (n=5 mice, One-way ANOVA (Tukey post-hoc), p values as stated). **C**.

Representative images of multiplex IHC staining of matrix proteins in wildtype and *Vil1-Grem1* mice, with quantification of changes in protein proportion between genotypes **D.** Representative images of multiplex IHC staining of immune cells in wildtype and *Vil1-Grem1* mice, with quantification of changes in cell proportion between genotypes. Scale bars 200µm.

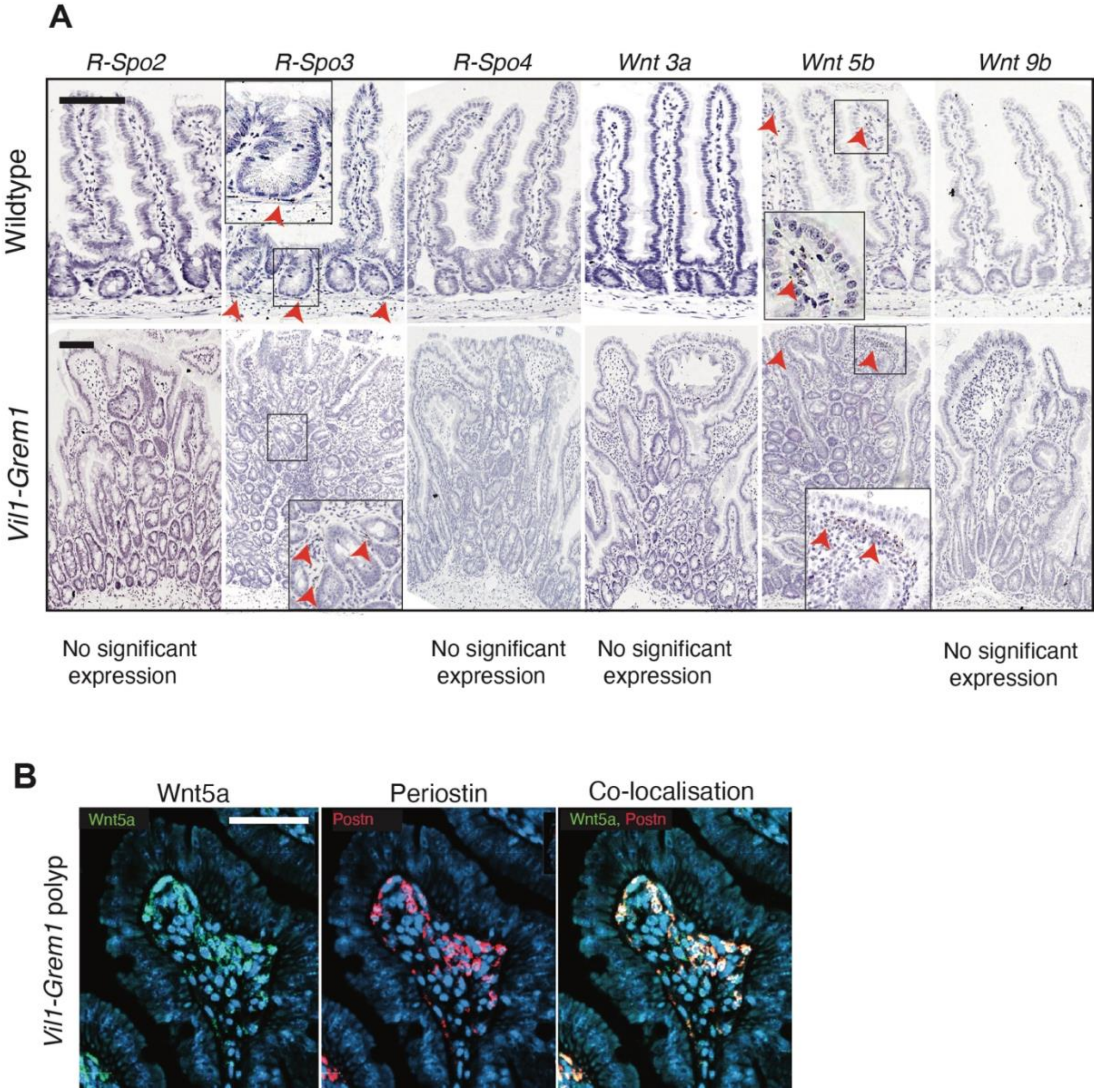


**Supplementary Figure 4. Additional Wnt ligand staining and co-staining. A.** Comparison of staining of additional Wnt ligands in wildtype and *Vil1-Grem1* mice, significantly expressed ligands identified with red arrows. **B.** Co-fluorescent ISH of *Wnt5a* and *Postn* at the periphery of *Vil1-Grem1* polyps. Scale bars 200µm.

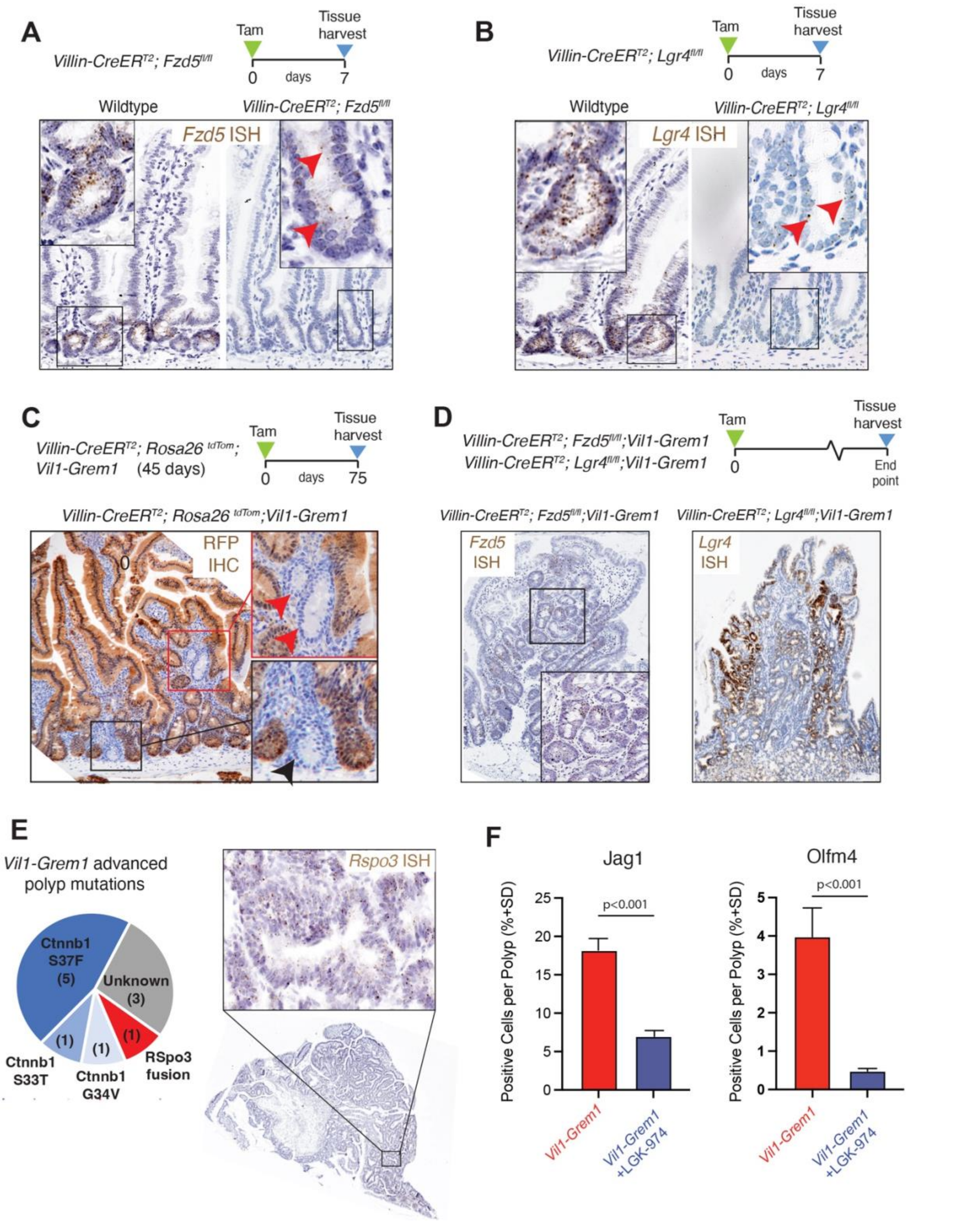


**Supplementary Figure 5. Wnt receptor knockout and advanced *Vil1-Grem1* polyp mutation burden.**

**A.** Schematic showing harvesting timepoint of *Villin-CreER*$^{T2}$,*Fzd5*$^{fl/fl}$ animals. *Fzd5* ISH staining (brown dots) of wildtype and *Villin-CreER*$^{T2}$*; Fzd5*$^{fl/fl}$ small intestine shows ongoing low level RNA expression post tamoxifen recombination in a small number of cells in the normal crypt base (red arrowheads).

**B.** Schematic showing harvesting timepoint of *Villin-CreER$^{T2}$; Lgr4$^{fl/fl}$ animals*. *Lgr4* ISH staining (brown dots) of wildtype and *Villin-CreER$^{T2}$; Lgr4$^{fl/fl}$* small intestine shows ongoing low level RNA expression post tamoxifen recombination in a small number of cells in the normal crypt base (red arrowheads). **C.** Schematic showing harvesting timepoint in 120 day old *Villin-CreER$^{T2}$; Rosa26$^{tdTom}$; Vil1-Grem1* animals. The emergence of non-recombined (unstained) escaper crypts in both the crypt base (black arrowhead) and ectopic crypts (red arrowheads) is seen as early as 4 days after tamoxifen recombination. **D.** Schematic showing harvesting timepoints in *Villin-CreER$^{T2}$; Fzd5$^{fl/fl}$; Vil1-Grem1* and *Villin-CreER$^{T2}$; Lgr4$^{fl/fl}$; Vil1-Grem1* animals, recombined at weaning and aged to end point. Representative ISH staining from these mice show robust expression of *Fzd5* and *Lgr4* in established polyps, despite receptor knockout, as a consequence of selection of escaper crypts in developing polyps. **E.** Pie chart shows identified mutations in *Ctnnb1* (β-catenin), detected by Sanger sequencing from micro-dissected advanced (*Lgr5(+))* polyps in *Vil1-Grem1* animals. ISH confirms epithelial staining of *RSpo3* in a single polyp with an identified *Ptprk-Rspo3* fusion mutation. **F.** Quantification of staining shows reduction in *Jag1* and OLFM4 positive cells in *Vil1-Grem1* animal polyps following treatment with LGK-974 porcupine inhibitor (n=5 mice per group, t test, p values as stated). Scale bars 200µm.

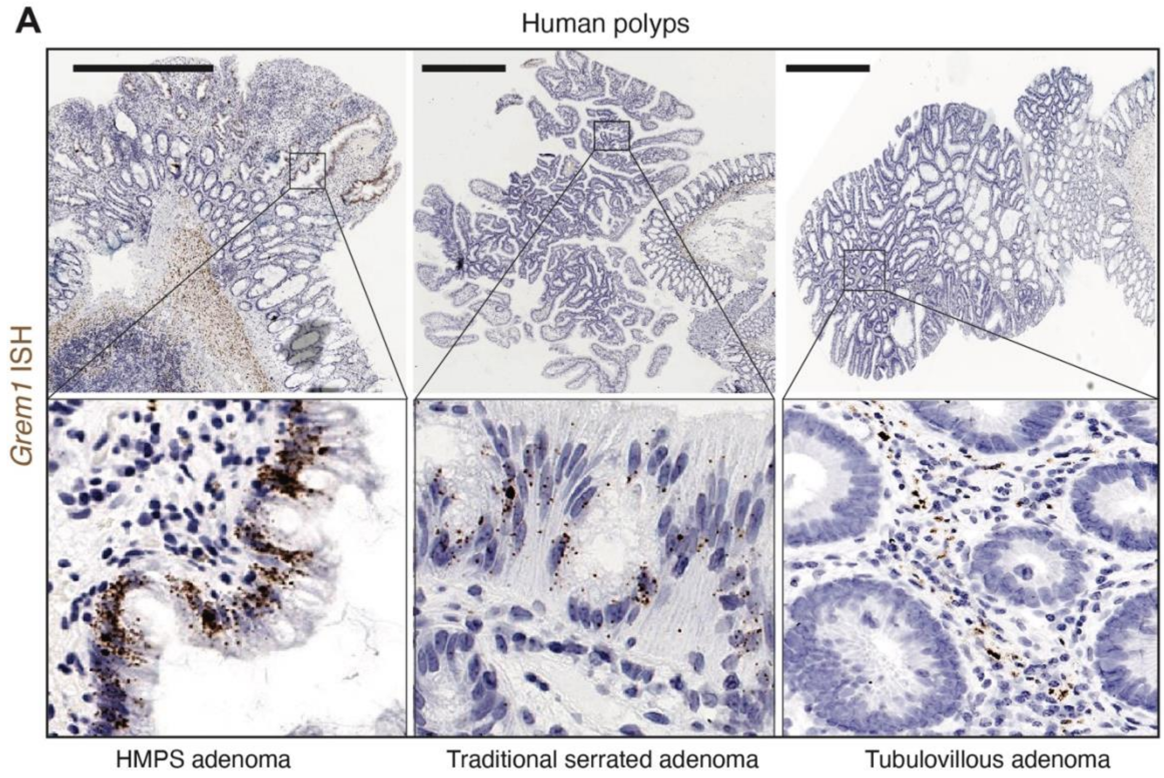


**Supplementary Figure 6. *GREM1* expression in HMPS and sporadic human polyps.** *GREM1* ISH (brown stain) in human polyps shows only stromal expression in sporadic tubulovillous adenomas (TVA) but marked epithelial expression in polyps from Hereditary Mixed Polyposis Syndrome (HMPS) and sporadic traditional serrated adenomas (TSA). Scale bars 500µm.
